# Out of the Mediterranean Region: worldwide biogeography of snapdragons and relatives (tribe Antirrhineae, Plantaginaceae)

**DOI:** 10.1101/855973

**Authors:** Juan Manuel Gorospe, David Monjas, Mario Fernández-Mazuecos

**Affiliations:** Real Jardín Botánico (RJB-CSIC), Plaza de Murillo 2, 28014 Madrid; Universidad Internacional Menéndez Pelayo (UIMP), Calle de Isaac Peral 23, 28040 Madrid

**Keywords:** Disjunction, Mediterranean Region, diversification rate, time-for-speciation effect, dispersal, species accumulation, proto-Mediterranean

## Abstract

**Aim:** The tribe Antirrhineae, including snapdragons, toadflaxes and relatives, is a model group for plant evolutionary research. It is widely distributed across the Northern Hemisphere and the Neotropics, but displays an uneven distribution of diversity, with more than 50% of species and subspecies in the Mediterranean Region. Here we conducted the first detailed, worldwide biogeographic analysis of the Antirrhineae and tested two alternative hypotheses (time-for-speciation vs. diversification rate differences) to explain the uneven distribution of diversity.

**Location:** Worldwide, with a focus on the Mediterranean Region.

**Taxon:** tribe Antirrhineae (Plantaginaceae).

**Methods:** A phylogenetic biogeographic approach was taken, accounting for area connections through time. Ancestral ranges, dispersal events, speciation and lineage accumulation within areas were estimated. Diversification rates for taxa present and absent in the Mediterranean Region were compared, accounting for the effect of a floral key innovation (nectar spur).

**Results:** A proto-Mediterranean origin in the Late Eocene was estimated, and the Mediterranean Region stood out as the main centre for speciation and dispersal. Congruent patterns of long-distance dispersal from the Mediterranean Region to North America were recovered for at least two amphiatlantic clades. A significant floristic exchange between the Mediterranean and south-western Asia was detected. We found no evidence of different diversification rates between lineages inside and outside the Mediterranean Region.

**Main conclusions:** The Mediterranean Region played a key role in the origin of the current distribution of the Antirrhineae. However, the higher species richness found in this region appears to be the result of a time-for-speciation effect rather than of increased diversification rates. The establishment of current mediterranean climates in the Northern Hemisphere appears to have contributed to the recent diversification of the group, in combination with colonisation of adjacent regions with arid and semi-arid climates.

## INTRODUCTION

Biogeographers have long been interested in patterns where similar organisms display distant distributions (disjunctions) and in the fact that some regions contain a large number of lineages while others are sparsely occupied (uneven distribution of diversity). Disjunct distributions have been a recurrent research topic in historical biogeography (Li, 1952; Raven, 1972; Sanmartín, Enghoff, & Ronquist, 2001; Thorne, 1972; Wen, 1999; Wen & Ickert-Bond, 2009). Even though they were historically explained by vicariance hypotheses, more recently the importance of long-distance dispersal has been highlighted (Sanmartín & Ronquist, 2004). Regarding the heterogeneity in the distribution of diversity, most studies have focused on the latitudinal gradient in species richness, while less attention has been paid to cases with an unbalanced distribution of richness between regions sharing similar latitude and climatic conditions (Valente, Savolainen, Manning, Goldblatt, & Vargas, 2011). Two main hypotheses have been proposed to explain the latter pattern (Wiens, 2011): (i) the time-for-speciation effect, which ascribes the higher species richness in some areas to a longer time for diversification (Mittelbach et al., 2007; Stephens & Wiens, 2002); and (ii) diversification rates differences, i.e. richness heterogeneity is better explained by spatial differences in rates of speciation and/or extinction (Mittelbach et al., 2007; Ricklefs, 2006). Differences in ecological limits to diversification have been proposed as a third hypothesis (Cook, Hardy, & Crisp, 2015; Rabosky, 2009), which can be considered a facet of diversification rate effects, and can only be properly distinguished when fossil evidence is available (Wiens, 2011). In any case, geography is regarded as one of the major determinants of diversification, in confluence with evolutionary innovations and environmental changes (Donoghue & Sanderson, 2015). Therefore, clades with discontinuous distribution patterns and uneven distribution of diversity become ideal systems to explore biogeographical and evolutionary processes.

The tribe Antirrhineae is a distinct clade of angiosperms within the Plantaginaceae family (Albach, Meudt, & Oxelman, 2005). It includes the common snapdragon (*Antirrhinum majus*), a well-known model species for plant biology (Schwarz-Sommer, Davies, & Hudson, 2003). As a result, the tribe as a whole can be considered a model group for evolutionary, ecological and developmental research (Cubas, Vincent, & Coen, 1999; Fernández-Mazuecos et al., 2019; Hileman, Kramer, & Baum, 2003). The Antirrhineae are characterized by having tubular flowers usually with a basal appendix or a spur (Sutton, 1988; Vargas, Rosselló, Oyama, & Güemes, 2004). The morphology of the seeds is very diverse and many dispersal strategies have been proposed, being anemochory and hydrochory the most recurrent ones (Elisens & Tomb, 1983; Sutton, 1988). Nevertheless, it has been indicated that seed traits do not seem specifically adapted to long-distance dispersal (Vargas et al., 2004). The tribe contains around 31 genera and more than 300 species (Fernández-Mazuecos et al., 2019; Sutton, 1988), and has an extremely limited fossil record mostly restricted to recent times (reviewed by Fernández-Mazuecos et al., 2019). At present, the Antirrhineae have a widespread distribution mostly in the Northern Hemisphere (from western North America in the West to Japan in the East), also extending to the Andean mountain ranges in the Southern Hemisphere (see map in Fig. 1a; Hong, 1983; Vargas et al., 2004). This distribution encompasses numerous climatic zones, the most representative ones being the two northern mediterranean climate regions (Mediterranean Basin and California) and their surroundings. In particular, the Mediterranean Basin (or Mediterranean Region) harbours more than half of the known representatives of the tribe (c. 53% of species and subspecies; Fig. 2a). Both northern mediterranean climate regions have been hypothesized as centres of diversification for the group (Sutton, 1988) but the processes responsible for the asymmetric distribution of species richness remain untested.

**Fig. 1.**
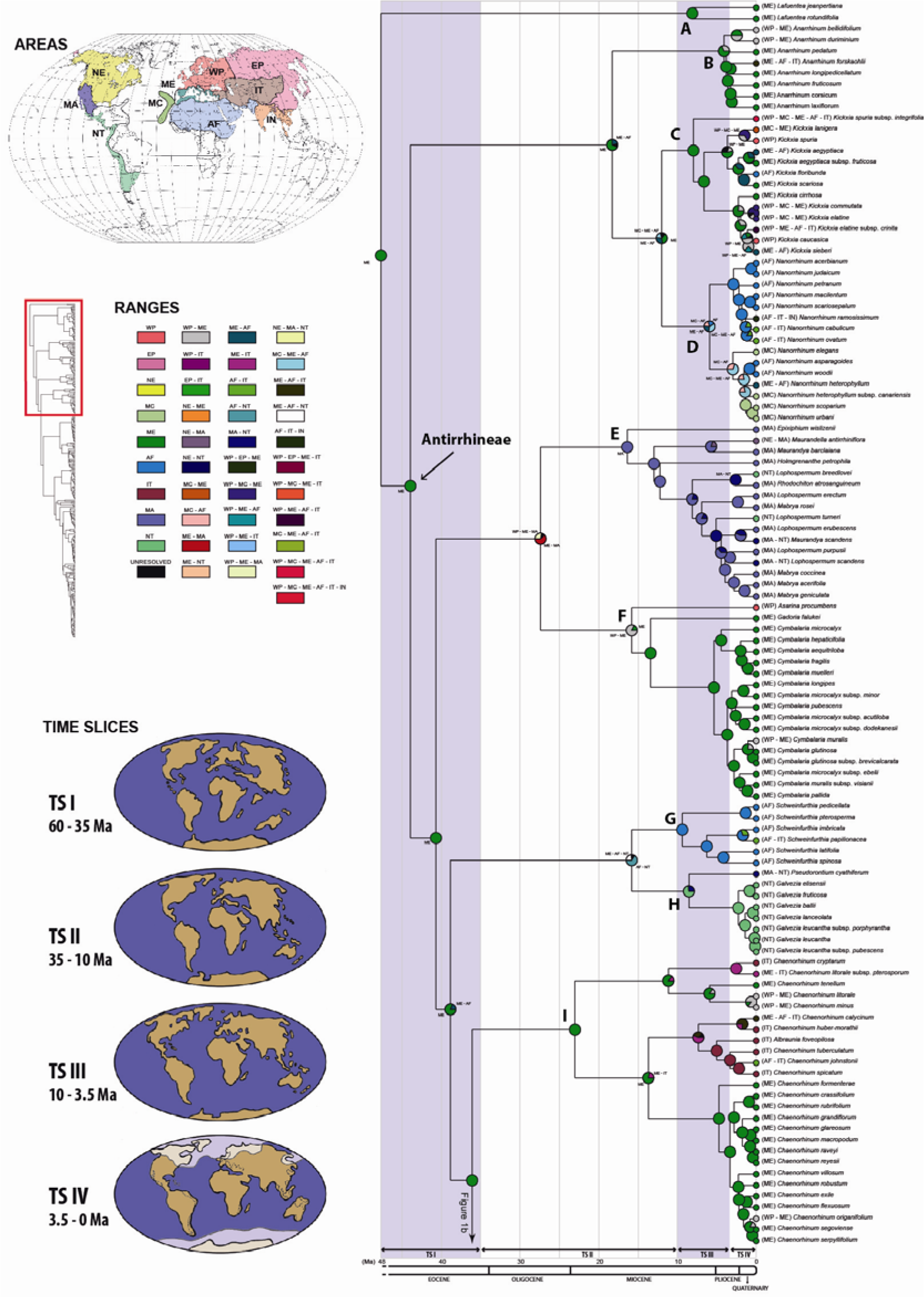
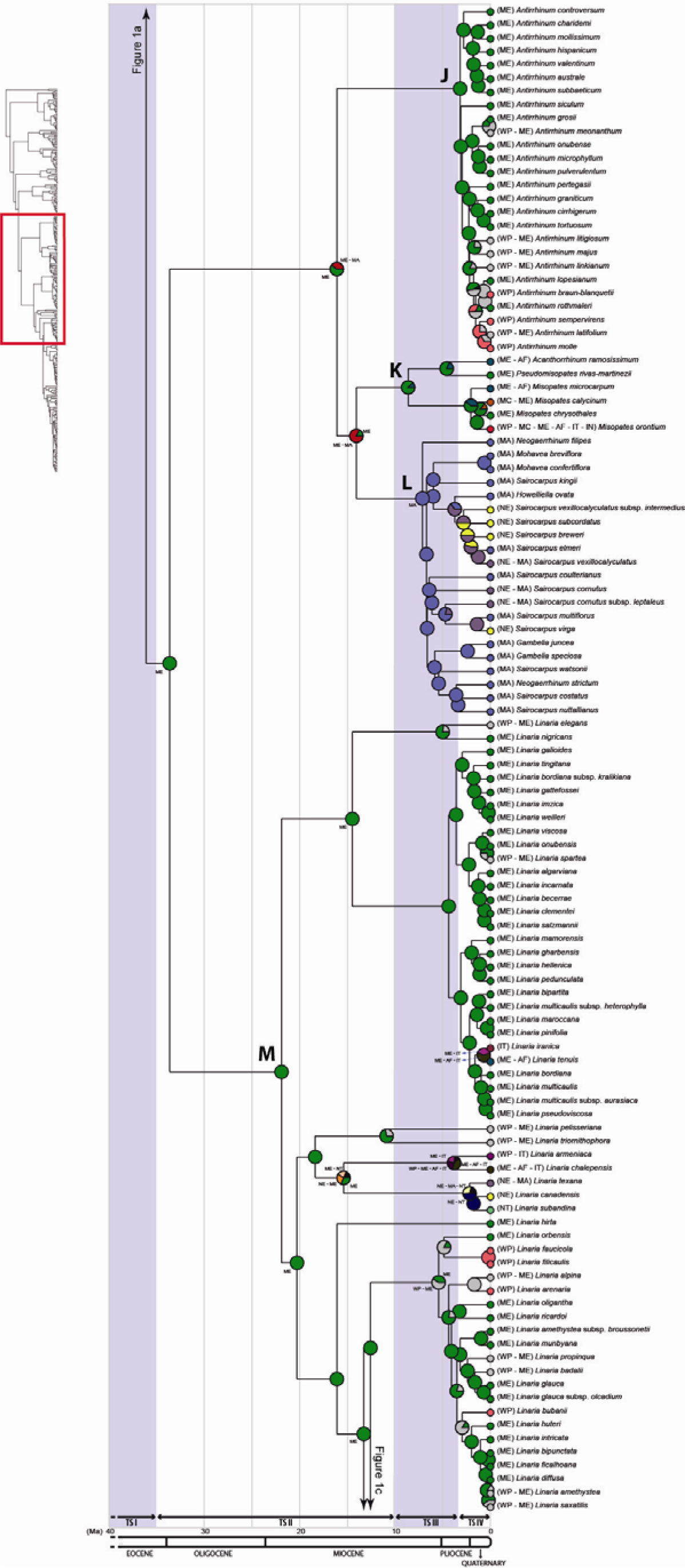
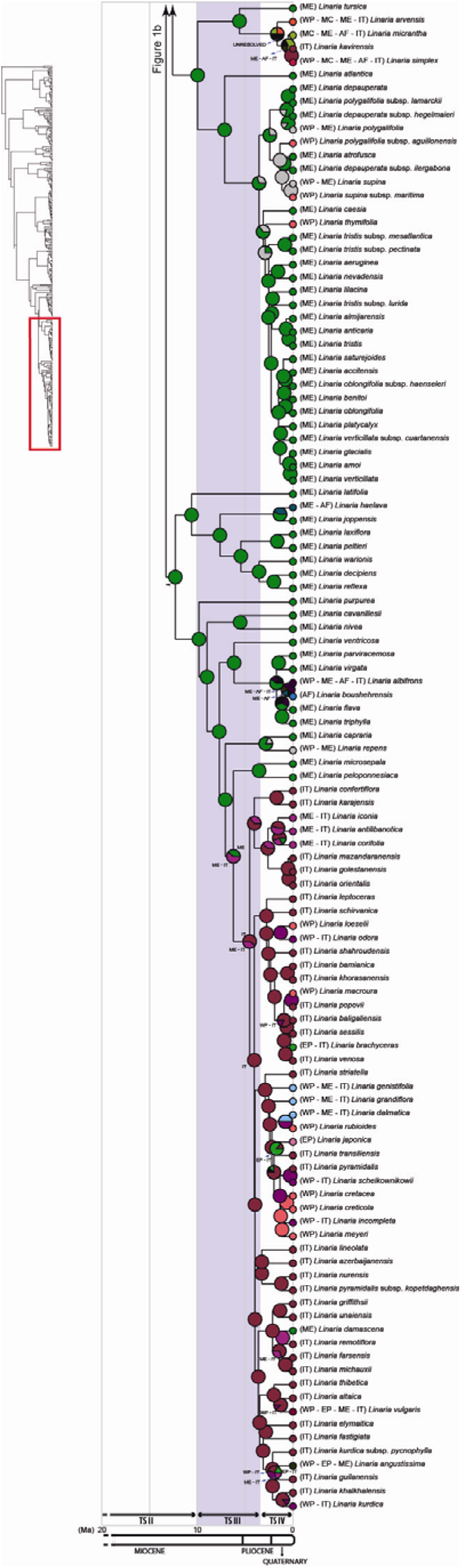
(a-c) Time-calibrated phylogeny of Antirrhineae and ancestral range estimation under the DEC model obtained in RASP. For simplicity only ancestral ranges (pie chart slices) with a probability over 0.15 are shown in colours, and at least two ranges were kept in all cases. Low-probability ranges are pooled in black. Letters at nodes (A to M) indicate major clades within the Anitrrhineae and the sister genus *Lafuentea* described in Appendix S1. Time slices (TSs) are denoted alternatively with blue and white backgrounds. The coloured area in the map at the top left in panel a represents the approximate distribution of the Antirrhineae. Maps representing a schematic configuration of landmasses for each TS (based on Meseguer et al., 2015) are shown at the bottom left in panel a. Areas: NE, Nearctic Region; WP, Western Palearctic Region; EP, Eastern Palearctic Region; MA, Madrean Region; ME, Mediterranean Region; IT, Irano-Turanian Region; NT, Neotropical Region; MC, Macaronesian Region; AF, African Region; IN, Indian-Indochinese Region. Time Slices: TSI, 60-35 Ma; TSII, 35-10 Ma; TSIII, 10-3.5 Ma; TSIV, 3.5-0 Ma.

**Fig. 2.**
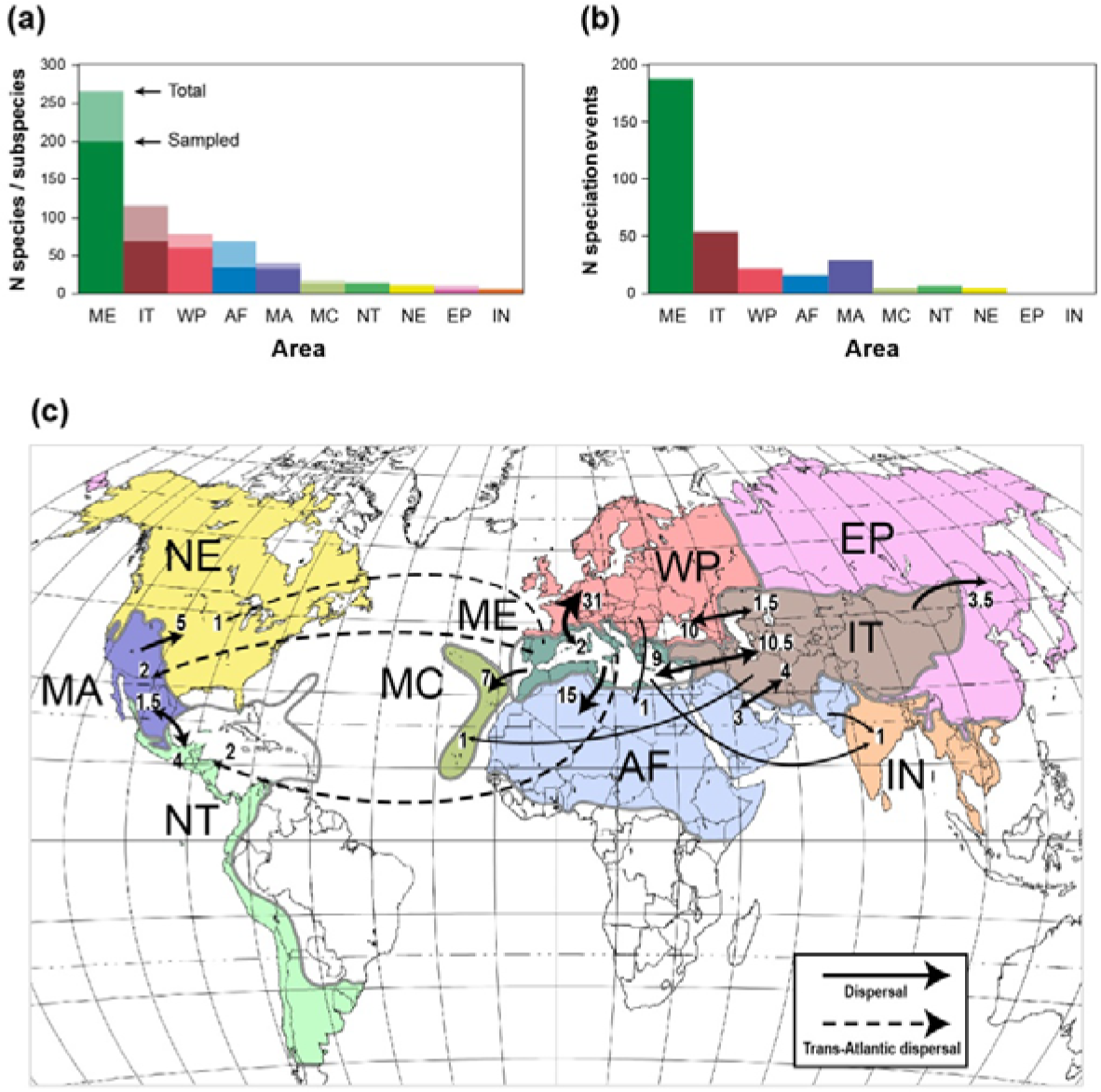
Species richness, speciation and dispersal events estimated for each area of the Antirrhineae distribution. (a) Number of sampled species and subspecies per area (dark coloured bars) in relation to the total number of known taxa according to a ‘splitter’ taxonomic treatment (light coloured bars). (b) Number of speciation events per area estimated under the DEC analysis in RASP. (c) Dispersal events between areas estimated under the DEC analysis in RASP. Arrow tip size is proportional to the number of dispersal events (numbers beside tips show the number of events between the connected areas in that direction), and dashed lines represent trans-Atlantic dispersal. Only events with values over 1 are represented. Areas: NE, Nearctic Region; WP, Western Palearctic Region; EP, Eastern Palearctic Region; MA, Madrean Region; ME, Mediterranean Region; IT, Irano-Turanian Region; NT, Neotropical Region; MC, Macaronesian Region; AF, African Region; IN, Indian-Indochinese Region.

The fact that some clades are distributed in both hypothetical centres of diversification has led to several hypotheses on the origin and events behind this amphiatlantic distribution. First, the Old World was suggested as the ancestral range for the tribe, due to the distribution of early-diverging genera revealed by phylogenetic reconstructions (Ghebrehiwet, Bremer, & Thulin, 2000; Vargas et al., 2004). Several colonization routes to the New World were proposed, including two terrestrial ones through land bridges at different times in the Cenozoic (the Thulean route and the Beringian route) or, alternatively, three independent events of long-distance colonization (Vargas et al., 2004). Later on, with the support of a genus-level, time-calibrated phylogeny and specific biogeographic analyses, Vargas et al. (2014) confirmed the Old World as the ancestral range and proposed four independent events of long-distance colonization from the Old World to the New World during the Miocene, tracing biogeographical congruence of four clades and ruling out vicariance scenarios (see also Ogutcen, Theriault, King, & Vamosi, 2017; Ogutcen & Vamosi, 2016). Additional biogeographic hypotheses have been proposed for specific groups within the tribe (Carnicero, Sáez, Garcia-Jacas, & Galbany-Casals, 2017; Elisens, 1985a, 1985b; Elisens & Nelson, 1993; Ghebrehiwet et al., 2000). Apart from these inferences, the precise ancestral range for the tribe and the detailed biogeographic events explaining its current worldwide distribution have not been explicitly assessed in a phylogenetic framework. The objectives of the present study were: (1) to estimate ancestral ranges for the Antirrhineae and its major clades based on a deeply sampled species-level phylogeny; (2) to test whether the Mediterranean Region was the centre of origin for the tribe; and (3) to compare diversification rates inside and outside the Mediterranean Region. The ultimate goal was to assess the causes of current diversity asymmetries by testing the time-for-speciation and diversification rate hypotheses.

## MATERIALS AND METHODS

### Data sampling and DNA sequencing

To incorporate available data in phylogenetic analyses, the supermatrix approach of Fernández-Mazuecos et al. (2019) was followed. We sampled 775 DNA sequences from 336 species and subspecies of Antirrhineae corresponding to the nuclear ribosomal internal transcribed spacers (ITS) and two plastid DNA (ptDNA) regions (*ndh*F and *rpl*32*-trn*L). Of these, 705 were retrieved from GenBank and 70 were newly generated (see Table S1 in Supporting Information for GenBank accession numbers, Table S2 for voucher specimens of newly sequenced taxa, and Appendix S1 for details on DNA extractions). Forty outgroup sequences representing 34 taxa and twelve families of Lamiales used by Fernández-Mazuecos et al. (2019) were also included. The three matrices (ITS, *ndh*F and *rpl*32-*trn*L) were then concatenated leading to a final dataset with 370 taxa and a total length of 3963 base pairs.

**Table 1.** Geography-dependent diversification in the Antirrhineae: binary approach (presence/absence in the Mediterranean Region) with hidden states. Log-likelihood (LnL) and Akaike information criterion (AIC) are shown for each model tested with the ‘hisse’ package under three alternative taxonomic treatments (‘splitter’, ‘intermediate’, ‘lumper’). Four models were tested: a character-independent model with two hidden states (CID-2), a character-independent model with four hidden states (CID-4), a character-dependent model without hidden states (BiSSE) and a character-dependent model with two hidden states (HiSSE). Models within 2 AIC units of the best model are shown in bold.

|  | Splitter |  |  | Intermediate |  |  | Lumper |  |  |
| --- | --- | --- | --- | --- | --- | --- | --- | --- | --- |
| Model | LnL | AIC | ΔAIC | LnL | AIC | ΔAIC | LnL | AIC | ΔAIC |
| CID-2 | -909.5 | 1843.1 | 15.5 | -837.3753 | 1698.751 | 16.915 | -741.8152 | 1507.63 | 25.609 |
| CID-4 | **-902.8** | **1827.6** | **0.0** | -832.5847 | 1687.169 | 5.333 | -732.7395 | 1487.479 | 5.458 |
| BiSSE | -929.1 | 1870.2 | 42.6 | -854.3531 | 1720.706 | 38.87 | -749.6548 | 1511.31 | 29.289 |
| HiSSE | **-898.5** | **1829.0** | **1.4** | **-824.9179** | **1681.836** | **0** | **-725.0105** | **1482.021** | **0** |

**Table 2.** State-dependent diversification: multi-state approach incorporating geography (presence/absence in the Mediterranean Region) and a floral key innovation (presence/absence of nectar spur). Log-likelihood (LnL) and Akaike information criterion (AIC) are shown for each diversification model tested with the ‘diversitree’ package under three alternative taxonomic treatments (‘splitter’, ‘intermediate’, ‘lumper’). The best model is shown in bold. λ diversification rate; µ, extinction rate; q, character transition rate.

|  | Splitter |  |  | Intermediate |  |  | Lumper |  |  |
| --- | --- | --- | --- | --- | --- | --- | --- | --- | --- |
| Model | LnL | AIC | ΔAIC | LnL | AIC | ΔAIC | LnL | AIC | ΔAIC |
| Full MuSSE | -932.4 | 1904.8 | 2.5 | -857.0 | 1754.1 | 2.3 | -753.6 | 1547.2 | 2.9 |
| Equal $\lambda$, $\mu$, $q$ | -988.7 | 1983.5 | 81.2 | -909.5 | 1825.0 | 73.2 | -795.4 | 1596.8 | 52.5 |
| Equal $\mu$, $q$ | -974.5 | 1961.0 | 58.7 | -894.3 | 1800.6 | 48.8 | -788.2 | 1588.3 | 44.1 |
| Equal $\lambda$, $q$ | -982.3 | 1976.6 | 74.4 | -903.4 | 1818.8 | 67.0 | -792.8 | 1597.5 | 53.3 |
| Equal $\lambda$, $\mu$ | -948.2 | 1924.4 | 22.2 | -873.1 | 1774.2 | 22.4 | -763.5 | 1555.0 | 10.7 |
| Equal $\lambda$ | -938.4 | 1910.8 | 8.5 | -863.6 | 1761.2 | 9.3 | -756.5 | 1547.1 | 2.8 |
| Equal $\mu$ | **-934.1** | **1902.3** | **0.0** | **-858.9** | **1751.8** | **0.0** | **-755.1** | **1544.3** | **0.0** |
| Equal $q$ | -972.0 | 1962.1 | 59.8 | -891.9 | 1801.9 | 50.1 | -785.7 | 1589.5 | 45.2 |

### Phylogenetic analyses and dating

The best-fitting substitution model for each DNA region was determined under the Akaike Information Criterion (AIC) in jModelTest 2.1.6 (Darriba, Taboada, Doallo, & Posada, 2012) using the CIPRES Science Gateway 3.3 (Miller, Pfeiffer, & Schwartz, 2010) (GTR+I+G for ITS and *ndh*F and GTR+G for *rpl*32*-trn*L). A preliminary topology was constructed with MrBayes 3.2.6 (Ronquist et al., 2012) in CIPRES using two runs with four chains, each with 10 million generations and a sampling frequency of 1000.

A time-calibrated phylogenetic analysis was performed in BEAST 2.4.2 (Bouckaert et al., 2014) using CIPRES with unlinked site models across partitions (as defined by jModelTest), unlinked clock models (uncorrelated relaxed clock in all cases), five MCMC chains and 250 million generations, sampling every 25000^th^ generation. Priors specified by Fernández-Mazuecos et al. (2019) were reproduced, including seven fossil calibrations (two within the Antirrhineae and five outside) and one secondary calibration (see Appendix S1 for further details). Chains were combined with LogCombiner 2.4.8 (included in BEAST 2 package), and a specific burn-in was removed from each chain based on trace plots visualised in Tracer 1.6 (Rambaut, Drummond, Xie, Baele, & Suchard, 2018). An effective sample size over 200 was obtained for all parameters, and a maximum clade credibility (MCC) tree representing common ancestor heights was generated with TreeAnnotator 1.8.4 (Drummond, Suchard, Xie, & Rambaut, 2012). Outgroup taxa, except *Lafuentea* (the sister genus to Antirrhineae), were pruned from the MCC tree. A final tree with 336 taxa was used in subsequent analyses.

### Biogeographic analyses

Ancestral range estimation was conducted using two methods implemented in the software RASP 4.0 (Yu, Harris, Blair, & He, 2015). The first one, Statistical Dispersal-Vicariance Analysis (S-DIVA) (Yu, Harris, & He, 2010), is a statistical version of the parsimony based Dispersal-Vicariance Analysis (DIVA) (Ronquist, 1997) and the second one takes a maximum likelihood (ML) approach under the Dispersal-Extinction-Cladogenesis (DEC) model (Ree, Moore, Webb, Donoghue, & Crandall, 2005; Ree & Smith, 2008).

Ten discrete areas were defined based on the floristic regions proposed by Takhtajan (1986) and the current known distribution of the Antirrhineae (Fig. 1a): Nearctic Region (NE); Madrean Region, including the mediterranean-climate region of California (MA); Neotropical Region (NT); Macaronesian Region (MC); Mediterranean Region (ME); African Region (AF); Western Palearctic Region (WP); Eastern Palearctic Region (EP); Irano-Turanian Region (IT); and Indian-Indochinese Region (IN) (see Appendix S1 for details). Distribution ranges for all terminal taxa were determined using taxonomic and floristic literature (see Table S3 for references). To improve computational performance ancestral ranges were constrained to a maximum of four areas in the DEC analysis (maximum range of 99% of sampled taxa). In the S-DIVA analysis, the widespread distribution of some terminal taxa led to estimates of widespread ancestors with high uncertainty at some nodes, which seems to be a tendency of DIVA methods (Buerki et al., 2011; Kodandaramaiah, 2010). Therefore, the number of areas was constrained to a maximum of three for the S-DIVA analysis (maximum range of 98% of sampled taxa).

In the DEC analysis, we accounted for changes in area connectivity through time caused by major tectonic events by following the paleogeographic model approach of Meseguer, Lobo, Ree, Beerling, and Sanmartín (2015). Information on paleogeographic changes was incorporated by scaling dispersal probability values between pairs of areas in four time slices (60-35 Ma, 35-10 Ma, 10-3.5 Ma and 3.5-0 Ma; see Appendix S1 and Fig. S1 for details). Dispersal events estimated under the DEC model were compiled in a map to illustrate the biogeographic connections between areas (values under one were not considered).

For comparison, we conducted ML biogeographic estimations in BioGeoBEARS (Matzke, 2013), with a maximum of four areas for ancestral and tip ranges to improve computational efficiency. Three biogeographical models were tested: DEC, DIVA-like and BayArea (Landis, Matzke, Moore, & Huelsenbeck, 2013). Models including jump dispersal were excluded because of the conceptual and statistical problems reported by Ree and Sanmartín (2018). The best-fitting model was chosen based on AIC values. Analyses were run with and without dispersal scalars (see above). To visualise geographical variation in the tempo of lineage accumulation, we generated lineages through space and time (LTST) plots for all areas using the R package ‘ltstR’ (Skeels, 2019). Plots were based on 10 biogeographic stochastic maps generated in BioGeoBEARS using the best-fitting biogeographic model (DEC) without dispersal scalars (due to the input requirements of ltstR).

### Geography-dependent diversification

We evaluated the potential effect of geography on rates of diversification inside and outside the Mediterranean Region using the hidden state speciation and extinction (HiSSE) method (Beaulieu & O’Meara, 2016). This is an extension of the binary state speciation and extinction (BiSSE) method (FitzJohn, Maddison, & Otto, 2009) that accounts for unmeasured factors (hidden traits) that may influence diversification rates in addition to the trait of interest. Two states were defined: presence in the Mediterranean Region (1), including Mediterranean endemics and widespread taxa also present in other regions; and absence from the Mediterranean Region (0). Four models where tested using the R package ‘hisse’ (Beaulieu & O’Meara, 2016): a character-independent model with two hidden states (CID-2); a character-independent model with four hidden states (CID-4); a character-dependent model without hidden states (BiSSE) and a character-dependent model with two hidden states (HiSSE). To account for taxonomic uncertainty, analyses were run under three alternative taxonomic treatments (‘splitter’, ‘intermediate’ and ‘lumper’) following the approach of Fernández-Mazuecos et al. (2019). Incomplete sampling was accounted for by incorporating state-specific sampling fractions in all models (Fig. 2a). After fitting the models, a marginal reconstruction of ancestral states and diversification rates for each model was performed. To integrate the uncertainty in model selection, all model reconstructions were averaged using AIC weights. The presence or absence in the Mediterranean Region was inferred for each node based on marginal probabilities (values over 0.5 were considered as presences in ME). Differences in diversification rates between states at nodes and tips were assessed using beanplots (Kampstra, 2008). The geography-specific method (GeoHiSSE) included in the package ‘hisse’, where an additional state for widespread taxa is incorporated, was discarded because the widespread state would group and treat very different distributions as equal. In fact, exploratory analyses with this method led to reconstructions that were inconsistent with those obtained using other methods.

Finally, we assessed the potential interaction between geography and a floral key innovation (nectar spurs; Fernández-Mazuecos et al., 2019) as drivers of diversification, as proposed by the ‘confluence’ hypothesis (Donoghue & Sanderson, 2015). To that end, we applied the multi-state speciation and extinction (MuSSE) model (FitzJohn, 2012) defining four states: absent from ME, without nectar spur (1); present in ME, without nectar spur (2); absent from ME, with nectar spur (3); and present in ME, with nectar spur (4). Analyses were run under the three alternative taxonomic treatments using the R package ‘diversitree’ (FitzJohn, 2012) accounting for incomplete sampling (Fig. 2a). A model with state-dependent speciation (λ) and extinction (μ) and asymmetrical transition rates (q) was compared against nested models with speciation rate, extinction rate and transition rate parameters constrained to be equal for all states. ML parameter values were calculated for each model, and models were compared based on AIC values. To obtain an estimate of parameter uncertainty, the best-fitting model was additionally explored using Bayesian inference, with exponentially distributed priors based on ML values. Each MCMC comprised 10000 steps, of which the first 1000 were discarded as burn-in.

## RESULTS

### Phylogenetic analyses and dating

The Antirrhineae were strongly supported as a monophyletic group (posterior probability, PP = 1) by both MrBayes and BEAST analyses (see Figs. S2, S3). The tribe was estimated to have diverged from the sister genus *Lafuentea* (A) during the Eocene, about 40–55 Ma (95% High Posterior Density, HPD), and an Eocene age was also estimated for the Most Recent Common Ancestor (MRCA) of the tribe, around 36–51 Ma (95% HPD). Major clades within the tribe (B to M; see Appendix S1 for details) were consistent with previous studies (Fernández-Mazuecos et al., 2019) and received strong support (PP ≈ 1) in both phylogenetic analyses, with the exception of clades J, K, L and M which obtained a lower support (PP = 0.8) in the BEAST analysis.

### Biogeographic analyses

S-DIVA and DEC analyses revealed similar ancestral range estimates for the root node and all major clades (Figs. 1a-c, S4). Both methods estimated the Mediterranean Region as the ancestral range for the Antirrhineae. This region was also estimated as the Most Likely Ancestral Range (MLAR) for most of the main clades in the tribe (B, C, I, J, K and M). Three major clades were estimated to have an American origin: two (E and L) in the Madrean Region and one (H) in the Neotropics. A widespread MLAR in WP-ME was estimated for clade F, while an origin in the African Region was estimated for clade G. The results were inconclusive for clade D, where ranges composed by different combinations of MC, ME and AF were obtained (Figs. 1a, S4).

The Mediterranean Region was the main area of origin for dispersal events, with over half of the total estimated events (around 69) (Fig. 2c). Adjacent areas WP and AF stood out as those most frequently colonized from ME (31 and 15 events respectively). Low levels of back-colonization of ME were estimated from most areas, with the exception of IT were colonization and back-colonization reached similar values (about 10 events). Colonization of Macaronesia from ME was also estimated (7 events). Dispersal across the Atlantic had a common origin in ME, and reached all three areas in the American continent (NE, MA and NT). Results show five events, but two of them occur along the same branch (alternatively from ME to NE or from ME to NT) leaving four trans-Atlantic dispersal events in total. Dispersal within the American continent was estimated mainly form MA to the surrounding areas NE and NT (5 and 4 events respectively). In Asia, colonization of EP was estimated to have an IT origin and no events of reverse colonization were estimated. Speciation was highest in ME (188 events). IT was another important area for speciation in the Old World (54 events), while MA was the main centre for speciation in the New World (29 events; see Fig. 2b).

Model testing in BioGeoBEARS revealed DEC as the best-fitting model (Table S4). Ancestral range estimation under DEC in BioGeoBEARS showed higher uncertainty for some nodes (Fig. S5) but it was generally consistent with DEC and S-DIVA analyses in RASP. Lineages through space and time (LTST) plots revealed a continued accumulation of lineages in ME since the Eocene (c. 40 Ma). In other regions, lineage accumulation started more recently after colonisation from ME. A steep increase in lineage accumulation in the Pliocene-Pleistocene (<5 Ma) was detected not only for ME, but also for IT, WP and AF. In contrast, in MA the fastest accumulation was inferred in the Late Miocene (5–10 Ma), followed by a slowdown in the Pliocene-Pleistocene (Fig. 3).

**Fig. 3.**
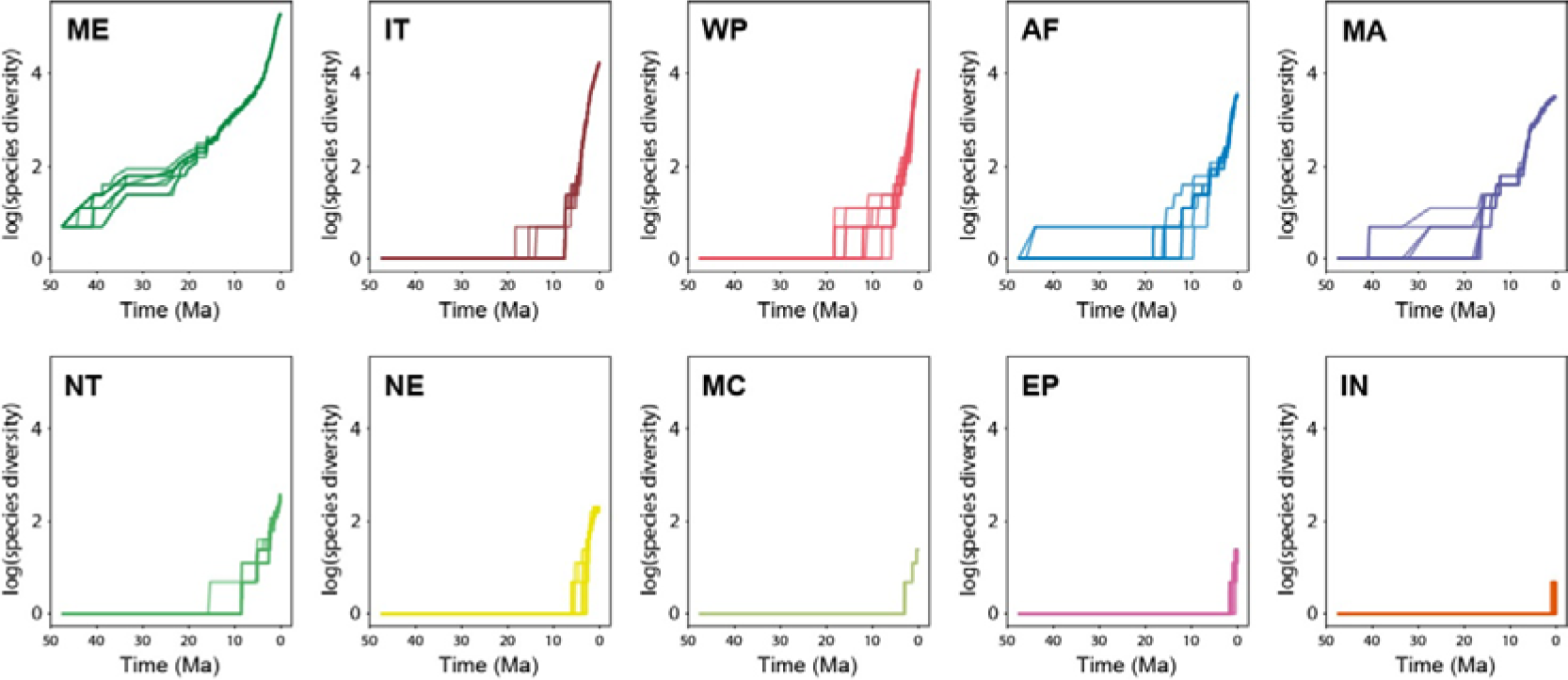
Accumulation of Antirrhineae lineages through time in each area (LTST plots) obtained from 10 biogeographic stochastic maps generated in BioGeoBEARS. Areas: NE, Nearctic Region; WP, Western Palearctic Region; EP, Eastern Palearctic Region; MA, Madrean Region; ME, Mediterranean Region; IT, Irano-Turanian Region; NT, Neotropical Region; MC, Macaronesian Region; AF, African Region; IN, Indian-Indochinese Region.

### Geography-dependent diversification

Analyses evaluating the effect of geography on net diversification showed a higher support for the full HiSSE model under the intermediate and lumper taxonomic treatments. Under the splitter treatment, CID-4 and HiSSE showed similar support (ΔAIC<2; Table 1). The model-averaged marginal reconstruction was consistent with other biogeographic analyses (Fig. 4a). The bean plot representation varied between taxonomic treatments, however the mean net diversification was similar (around 0.2) between states and taxonomic treatments (Fig. 4b).

**Fig. 4.**
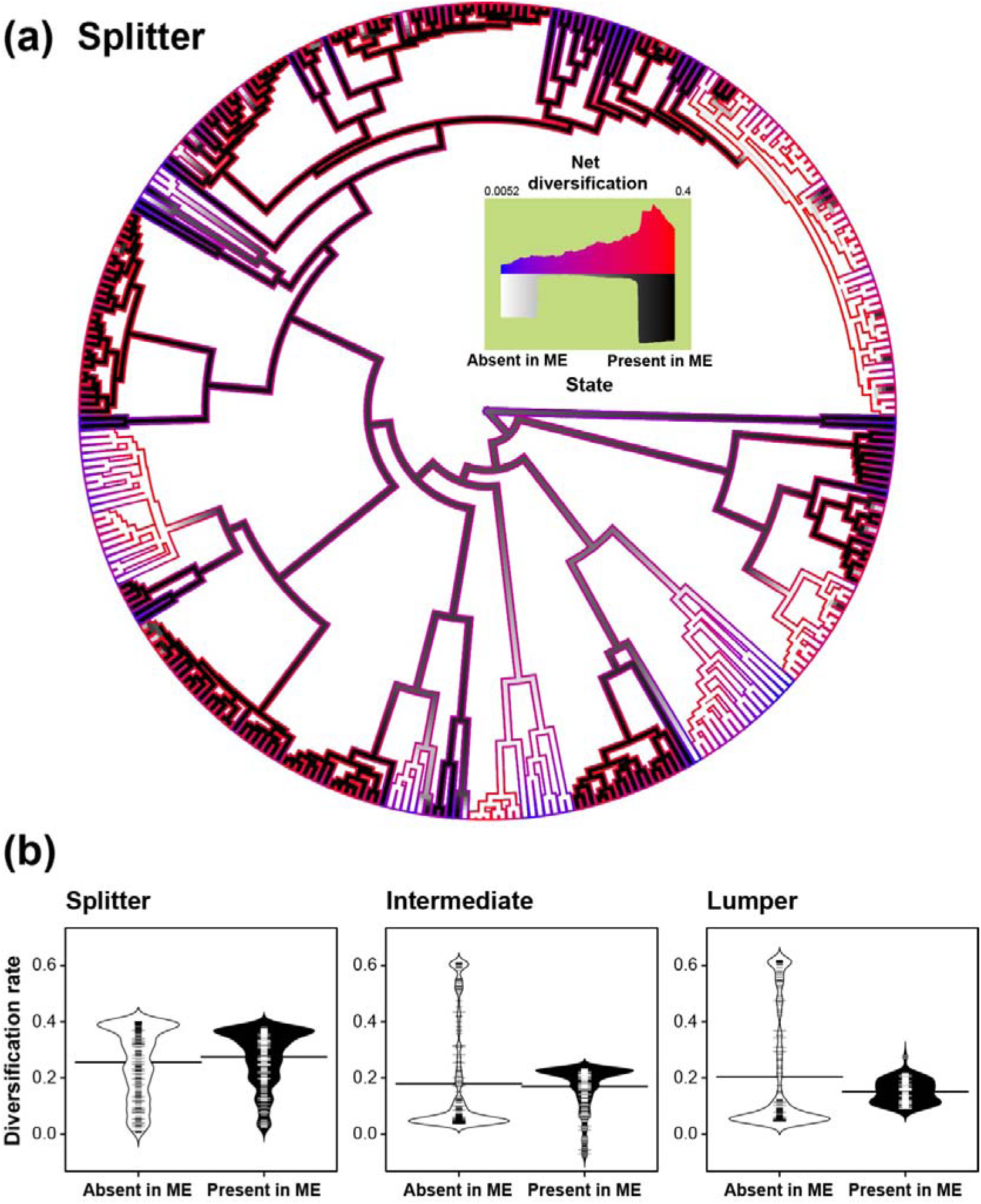
Effect of geography (presence/absence in the Mediterranean Region) on diversification of the Antirrhineae, as estimated by the ‘hisse’ package. (a) Model-averaged marginal reconstruction under the ‘splitter’ taxonomic treatment; four models were averaged: a character-independent model with two hidden states (CID-2), a character-independent model with four hidden states (CID-4), a character-dependent model without hidden states (BiSSE) and a character-dependent model with two hidden states (HiSSE); net diversification rates are represented as colour shading along branch edges and presence/absence in the Mediterranean is represented in black and white respectively. (b) Beanplots representing the distribution of net diversification rates at nodes and tips estimated after averaging four models (CID-2, CID-4, BiSSE and HiSSE) under three alternative taxonomic treatments (‘splitter’, ‘intermediate’, ‘lumper’); horizontal bars indicate mean values.

In MuSSE analyses, the equal extinction model (Equal μ) received the strongest support under all taxonomic treatments (Table 2). Bayesian posterior distributions of diversification rates under the equal extinction model showed similar results for all taxonomic treatments. The non-Mediterranean, spurred state showed higher diversification rate estimates (around 0.4) than the rest of states, with low overlap of 95% HPD intervals. Spurred taxa present in ME received intermediate diversification values (around 0.2) and spurless taxa (both present and absent in ME) had the lowest estimates (around 0.1; Fig. 5).

**Fig. 5.**
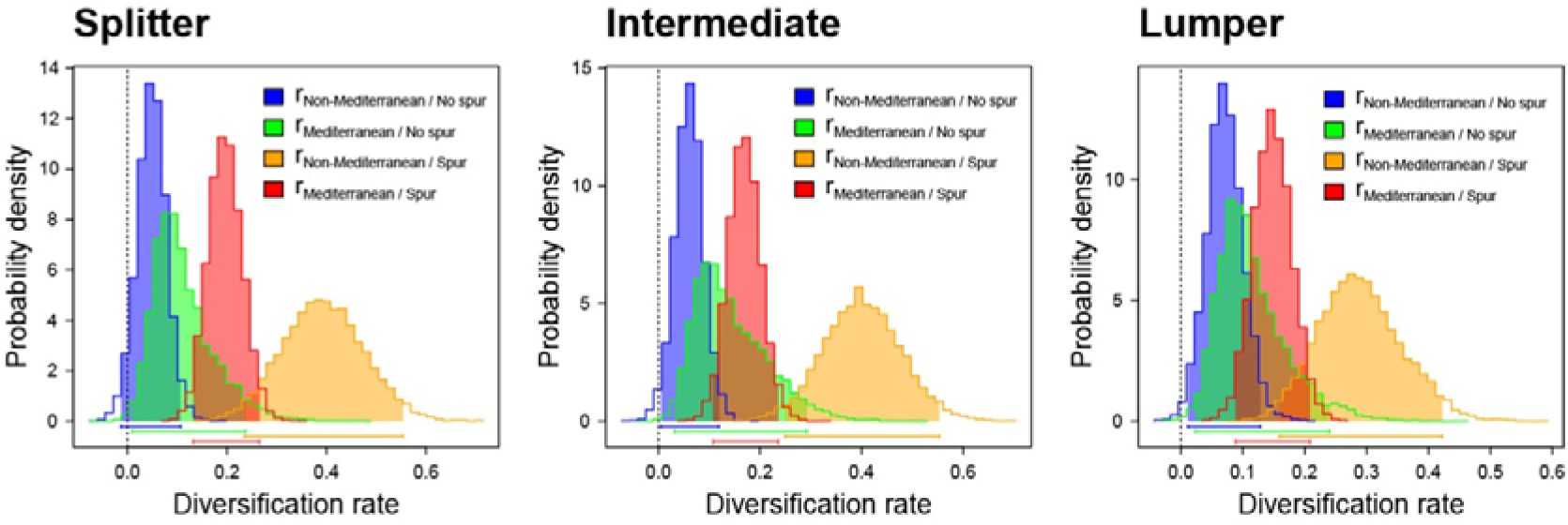
Interaction between geography (presence/absence in the Mediterranean Region) and a floral key innovation (presence/absence of nectar spur) as drivers of diversification of the Antirrhineae. Bayesian posterior distribution of diversification rates for the equal extinction model (Equal µ) under three alternative taxonomic treatments (‘splitter’, ‘intermediate’, ‘lumper’). r, rates of diversification.

## DISCUSSION

The present study provides the first worldwide analysis of biogeographical patterns in the tribe Antirrhineae based on a deep taxonomic and geographical sampling. Results were consistent across analyses and with previous phylogenetic studies, allowing us to test with a relatively high certainty the influence of geography in the evolutionary history of this group.

### Antirrhineae origin and diversification

Ancestral range estimations revealed a proto-Mediterranean origin for the clade during the Late Eocene (Fig. 1a), thus confirming our hypothesis and supporting previous suggestions (Vargas et al., 2004; Vargas et al., 2014). In the Eocene (55.8–33.9 Ma), the northward drift of the Indian and Australian plates led to the closure of the Tethys Ocean, leaving a proto-Mediterranean Sea in the West, between the African-Arabian and Eurasian plates (see Fig. 1a, time slice I; Meulenkamp & Sissingh, 2003; Rögl, 1999). The gradual climatic cooling during the Eocene (Zachos, Dickens, & Zeebe, 2008) favoured the development of a mixed-mesophytic forest around the Eocene-Oligocene boundary in northern mid-latitudes (Meseguer et al., 2015; Morley, 2007; Wolfe, 1975). We estimated that the common ancestor of the Antirrhineae occurred during this period (Fig. 1a), pointing out the possibility of a connection between the tribe’s differentiation and vegetational changes driven by climatic conditions and geographical events.

Around the mid-Miocene, the collision between the African-Arabian and Eurasian plates originated the current configuration of the Mediterranean Basin (Jolivet, Augier, Robin, Suc, & Rouchy, 2006). During the Miocene, aridity increased (Griffin, 2002) and a sclerophyllous ancestral flora was established in this region (Thompson, 2005), where ancestors of major Antirrhineae clades might have undergone diversification (Fig. 1a, b) followed by expansion to other areas (Fig. 3). This differentiation of major clades, prior to the onset of the mediterranean climate, has also been detected for several Mediterranean plant lineages (Vargas, Fernández-Mazuecos, & Heleno, 2018). The relatively recent diversification within most Antirrhineae clades is consistent with the appearance of mediterranean climates during the Pliocene in the Mediterranean Basin and during the Late Miocene-Pliocene in south-western North America (Millar, 2012; Rundel et al., 2016; Suc, 1984) (Figs.2b, 3). The Madrean Region in south-western North America, including the mediterranean-climate region of California, seems to share a similar diversification and lineage expansion pattern with the Mediterranean Region, even though speciation and dispersal estimates were lower for the American continent, possibly as a result of a more recent arrival of the Antirrhineae (Figs. 2b, c, 3).

### Major dispersal routes and distributional patterns

The Mediterranean Region was inferred as the major centre of dispersal for the tribe, as shown by the numerous colonization events from this area to surrounding ones in all directions (Fig. 2c). In contrast, a low number of back-colonization events was detected from other areas to the Mediterranean Region, with the exception of the Irano-Turanian Region, with which an ongoing floristic exchange is apparent (see below).

Our results showed that long-distance dispersal is a relevant process to explain the current amphiatlantic distribution of the Antirrhineae, as already suggested by Vargas et al. (2014). Despite the fact that most Antirrhineae seeds do not show specific long-distance dispersal adaptations, it has been indicated that dispersal mechanisms unrelated to dispersal syndromes are often relevant means of long-distance colonization (Guzmán & Vargas, 2005, 2009b; Higgins, Nathan, & Cain, 2003; Vargas, Heleno, Traveset, & Nogales, 2012). The importance of dispersal across large distances is increasingly being considered in biogeography. Indeed, some transoceanic disjunctions, especially in plant groups, are better explained by long-distance dispersal than by vicariance when a phylogenetic biogeography approach is taken (Sanmartín et al., 2001; Sanmartín & Ronquist, 2004). Our deep sampling and detailed analyses allowed us to pinpoint specific colonization routes: a northern route for two of the four long-distance migrations across the Atlantic (from the Mediterranean Region to the Madrean Region) and a central Atlantic route (from either the Mediterranean or African regions to the Neotropical Region); the route for the remaining dispersal event is still uncertain (Fig. 2c). This connection between both sides of the Atlantic during the Miocene has also been estimated for other plant groups (Davis, Fritsch, Li, & Donoghue, 2002; Fiz et al., 2008; Guzmán & Vargas, 2009a; Vargas et al., 2014). Also, disjunct distributions of other plant taxa have been proposed as the outcome of trans-Atlantic long-distance dispersal (Kadereit & Baldwin, 2012).

### Floristic exchange between the Mediterranean and Irano-Turanian regions

Results showed the Irano-Turanian Region as an important area for diversification, especially for one of the three main clades of *Chaenorhinum* and a clade of *Linaria* formed by *L.* sect. *Linaria* and *L.* sect. *Speciosae* (Fig. 1a, c). *Linaria* sect. *Linaria* includes the few Antirrhineae taxa that reached the Eastern Palearctic Region through the Irano-Turanian Region (Figs. 1c, 2c). This group is characterized by having winged seeds (Sutton, 1988) which might have favoured dispersal. Speciation events within the Irano-Turanian Region are consistent with a recent diversification since the Pliocene from ancestors adapted to dry climates, which were already established in this area since the Late Miocene (Manafzadeh, Salvo, & Conti, 2014). We estimated a high rate of floristic exchange between the Irano-Turanian Region and the Mediterranean Region, highlighting the importance of this biogeographic connection (Fig. 2c). A continuous exchange of taxa between these areas was also obtained for subfamily Apioideae (Banasiak et al., 2013), and the Irano-Turanian Region was a colonization route for other plant groups (Emadzade, Gehrke, Linder, & Hörandl, 2011; Jabbour & Renner, 2012; Manafzadeh et al., 2014; Mansion et al., 2008).

### Influence of geography on diversification rates

Character-dependent or character-independent models were supported by HiSSE analyses depending on the taxonomic treatment (Table 1). Regardless, diversification rate values were not significantly different between states and treatments (Fig. 4b), indicating that the presence or absence in the Mediterranean Region does not have a significant impact on diversification within the group. Considering that the Mediterranean Region was estimated as the ancestral range for the tribe and for most major clades, the higher species richness found in the Mediterranean Region is best explained by the time-for-speciation hypothesis (Stephens & Wiens, 2002). Indeed, LTST plots detected a longer period of species accumulation in the Mediterranean Region (since the Eocene) than in other areas, where lineage accumulation started around the Miocene after colonisation form the Mediterranean (Fig. 3). Ancestral adaptation to dry conditions in the Mediterranean, developed during the Miocene-Pliocene, may have favoured colonization of and diversification in other arid and semi-arid regions of the Northern Hemisphere, such as the Madrean Region in western North America and the Irano-Turanian Region in south-western Asia. In contrast, the high accumulation of lineages detected in the Western Palearctic and African regions appears to be the result of a high number of colonization events (mostly from the Mediterranean) rather than of high diversification within these areas (Figs. 2c, 3).

As previous authors have suggested (Donoghue & Sanderson, 2015; Fernández-Mazuecos et al., 2019; Guzmán, Gómez, & Vargas, 2015) a multi-state approach that takes into consideration a combination of factors would help explain disparities in species richness. Our analysis of the interaction between geography and a floral key innovation as drivers of diversification again failed to show an increased diversification rate in the Mediterranean Region (Fig. 5), thus providing further support for the time-for-speciation hypothesis. Taxa with nectar spur generally showed higher diversification rates, coinciding with Fernández-Mazuecos et al. (2019), and results suggest a complex interaction with geography and possibly additional factors driving diversification. Interestingly, the higher diversification rate detected for the non-Mediterranean taxa with nectar spur is attributable to the recent expansion of a single *Linaria* clade (formed by *L.* sect. *Linaria* and *L.* sect. *Speciosae*) into the Irano-Turanian Region.

In conclusion, an origin for the tribe Antirrhineae was estimated in the ancient Mediterranean during the Late Eocene. The highest number of speciation events was estimated for the Mediterranean Region, which was also the centre of dispersal for the tribe. South-western Asia (the Irano-Turanian Region) was revealed as another relevant area in the evolutionary history of the Antirrhineae, with a high number of speciation events and a recurrent floristic exchange with the Mediterranean Region. The establishment of mediterranean climates in the Northern Hemisphere around the Pliocene seems to have played a key role in the recent diversification of the tribe. However, the presence of a higher number of taxa within the Mediterranean Region cannot be explained by higher diversification rates. Instead, it seems to be the result of species accumulation through time. The combination of evolutionary and ecological approaches through integrative historical biogeography would contribute to better understand the ensemble of biotic and abiotic factors that have driven diversification in the Antirrhineae.

## Data availability statement

New DNA sequences were deposited in the GenBank database (see Table S1 for accession numbers).

## ACKNOWLEDGEMENTS

The authors thank Emilio Cano and Olga Popova for laboratory assistance; and Isabel Sanmartín, Ana Otero, Pablo Vargas and Alex Skeels for their useful comments. This research was part of J.M.G.’s MSc thesis (Master in Biodiversity in Tropical Areas and Conservation, Menéndez Pelayo International University – CSIC). D.M. was supported by the Youth Employment Initiative (European Social Fundand Education, Youth and Sport Office, Community of Madrid; reference PEJD-2017-PRE/AMB-3612). M.F.-M. was supported by a Juan de la Cierva fellowship (Spanish Ministry of Economy and Competitiveness; reference IJCI-2015-23459).

## Biosketch

Juan Manuel Gorospe is a MSc researcher interested in plant biodiversity and conservation using geographic and evolutionary approaches. ORCID: https://orcid.org/0000-0002-8118-5785

David Monjas is a research and laboratory assistant centred in the study of the evolution of organisms.

Mario Fernández-Mazuecos is a postdoctoral researcher investigating plant biodiversity from evolutionary and ecological perspectives. ORCID: https://orcid.org/0000-0003-4027-6477

Author contributions: J.M.G. and M.F.-M. conceived the ideas; J.M.G. and D.M. collected the data; J.M.G. and M.F.-M. analysed the data; and J.M.G. led the writing.

## Supporting Information

**Appendix S1.** Supplementary text.

**Table S1.** GenBank accession numbers for previously published and newly generated DNA sequences of Antirrhineae and the outgroup used in the present study.

**Table S2.** Vouchers specimens for newly-sequenced taxa of Antirrhineae.

**Table S3.** Distribution ranges of taxa used in biogeographic analyses. In the cases where subspecies are included, the range under the species name represents that of the type subspecies. Sources of information are provided. Areas: NE, Nearctic Region; WP, Western Palearctic Region; EP, Eastern Palearctic Region; MA, Madrean Region; ME, Mediterranean Region; IT, Irano-Turanian Region; NT, Neotropical Region; MC, Macaronesian Region; AF, African Region; IN, Indian-Indochinese Region.

**Table S4.** Comparison of biogeographic models in BioGeoBEARS. Log-likelihood (LnL) and Akaike information criterion (AIC) are shown for each model. The best model is shown in bold.

**Fig. S1.** Dispersal probability matrices for each time slice (TS), and maps representing a schematic configuration of landmasses for each TS (based on Meseguer et al., 2015; see Appendix S1 for further information about dispersal probability values and time slices). Areas: NE, Nearctic Region; WP, Western Palearctic Region; EP, Eastern Palearctic Region; MA, Madrean Region; ME, Mediterranean Region; IT, Irano-Turanian Region; NT, Neotropical Region; MC, Macaronesian Region; AF, African Region; IN, Indian-Indochinese Region. Time Slices: TSI, 60-35 Ma; TSII, 35-10 Ma; TSIII, 10-3.5 Ma; TSIV, 3.5-0 Ma.

**Fig. S2.** Bayesian phylogenetic tree of Antirrhineae based on analysis of ITS, *ndh*F and *rpl*32-*trn*L sequences in MrBayes. Numbers above branches are Bayesian posterior probabilities. Node letters (A to M) indicate major clades within the Antirrhineae and the sister genus *Lafuentea* described in Appendix S1. The most recent common ancestors for the Plantaginaceae and the Antirrhineae are indicated and family names for outgroup taxa used for calibration are included.

**Fig. S3.** Maximum clade credibility tree from phylogenetic analysis in BEAST showing the Posterior Probability value for node support, blue bars indicate divergence time estimates at a 95% High Posterior Density. All other conventions as in Fig. S2.

**Fig. S4.** S-DIVA ancestral range estimation in RASP. Node letters (A to M) correspond to the major clades within Anitrrhineae and the sister genus *Lafuentea* described in Appendix S1. The most recent common ancestor for the Antirrhineae is indicated. The coloured range in the map represents the approximate distribution of the Antirrhineae. Areas: NE, Nearctic Region; WP, Western Palearctic Region; EP, Eastern Palearctic Region; MA, Madrean Region; ME, Mediterranean Region; IT, Irano-Turanian Region; NT, Neotropical Region; MC, Macaronesian Region; AF, African Region; IN, Indian-Indochinese Region.

**Fig. S5.** Ancestral range estimation under the DEC model in BioGeoBEARS (with dispersal scalars). Letters at nodes and branches correspond to the range with the highest probability. Areas: A, Western Palearctic Region; B, Eastern Palearctic Region; C, Nearctic Region; D, Madrean Region; E, Mediterranean Region; F, African Region; G, Irano-Turanian Region; H, Macaronesian Region; I, Indian-Indochinese Region; J, Neotropical Region.

