## Supporting Information for "Out of the Mediterranean Region: worldwide biogeography of snapdragons and relatives (tribe Antirrhineae, Plantaginaceae)"

**Appendix S1.** Supplementary text.

**Table S1.** GenBank accession numbers for previously published and newly generated DNA sequences of Antirrhineae and the outgroup used in the present study.

**Table S2.** Vouchers specimens for newly-sequenced taxa of Antirrhineae.

**Table S3.** Distribution ranges of taxa used in biogeographic analyses.

**Table S4.** Comparison of biogeographic models in BioGeoBEARS.

**Fig. S1.** Dispersal probability matrices for each time slice and maps representing a schematic configuration of landmasses.

**Fig. S2.** Bayesian phylogenetic tree of Antirrhineae based on analysis of ITS, *ndh*F and *rpl*32-*trn*L sequences in MrBayes.

**Fig. S3.** Maximum clade credibility tree from phylogenetic analysis in BEAST.

**Fig. S4.** S-DIVA ancestral range estimation in RASP.

**Fig. S5.** Ancestral range estimation under the DEC model in BioGeoBEARS (with dispersal scalars).

**Appendix S1.** Supplementary text.

### MATERIALS AND METHODS

#### *DNA sequencing*

DNA regions ITS, *ndh*F and *rpl*32-*trn*L were selected according to previous studies (Fernández-Mazuecos, Blanco-Pastor, Gómez, & Vargas, 2013; Fernández-Mazuecos et al., 2019; Fernández-Mazuecos, Blanco-Pastor, & Vargas, 2013; Fernández-Mazuecos & Vargas, 2011; Ghebrehiwet, Bremer, & Thulin, 2000; Vargas, Rosselló, Oyama, & Güemes, 2004).

DNA extractions were obtained using the DNeasy Plant Mini Kit (Qiagen, CA, USA). PCR amplification followed the methods of Fernández-Mazuecos, Blanco-Pastor, and Vargas (2013) for ITS and Fernández-Mazuecos and Vargas (2011) for *rpl*32-*trn*L. Amplified products were submitted to Macrogen Inc. (Macrogen Europe, Madrid, Spain) for Sanger sequencing using primers P1A and P4 for the ITS region (Sang, Crawford, & Stuessy, 1995; White, Bruns, Lee, & Taylor, 1990) and trnL(UAG) and rpl32F for the *rpl*32-*trn*L region (Shaw, Lickey, Schilling, & Small, 2007). Sequences were assembled in Geneious 5.1.7 (Kearse et al., 2012).

#### *Priors used for the time-calibrated phylogenetic analysis in BEAST*

*Plocosperma buxifolium* was established as the outgroup species following Schäferhoff et al. (2010) by constraining the remaining taxa as a monophyletic group. The age of this ingroup was secondarily calibrated by assigning a normal prior distribution to its Most Recent Common Ancestor (MRCA) with mean 74 Ma and standard deviation 1.25 Ma (Bell, Soltis, & Soltis, 2010). Two fossil calibrations within the Antirrhineae were included: fossil seeds form the Upper Pliocene identified as *Linaria vulgaris* (Dorofeev, 1963) were used to calibrate the stem age of the *Linaria* sect. *Linaria* + sect. *Speciosae* clade with a log-normal distribution (Offset = 2.6 Ma, S = 1.25 and M = 1.0) and fossil seeds from the Middle Miocene classified as *Asarina ruboidea* (Mai, 2001) were employed to calibrate the stem age of *Asarina* using a log-normal distribution (Offset = 11.6 Ma, S = 1.25 and M = 1.0). Based on the results of Blanco-Pastor, Vargas, and Pfeil (2012), *Linaria* sect. *Supinae* was set as monophyletic. Following Vargas et al. (2014), five more calibration points outside the Antirrhineae were implemented based on fossil taxa representing five Lamiales lineages (Oleaceae tribe Fraxineae, Bignoniaceae, Lamiaceae, Plantaginaceae tribe Gratioleae and Plantaginaceae tribe Plantagineae).

Two taxa (*Nanorrhinum sagittatum* and *Misopates salvagense*) were pruned from the final tree because morphology and phylogenetic results suggest they are conspecific with *Nanorrhinum heterophyllum* and *Misopates orontium* respectively.

*Delimitation of biogeographic regions*

Ten discrete areas were delimited based on the Antirrhineae's worldwide distribution and using Takhtajan's (1986) Floristic Regions of the World as reference. Details about areas considered, extents and limits are offered below:

<u>Western Palearctic Region (WP)</u>: It includes non-Mediterranean Europe, reaching the N-NW of the Iberian Peninsula (north-western Portugal, Galicia, Cantabrian Mountains and Pyrenees), the northern Apennines, northern Balkans, northern Anatolia and most of the Caucasus in the South and the Ural Mountains in the East.

Eastern Palearctic Region (EP): From the Ural Mountains in the West to the Japanese archipelago in the East, including all of Siberia (North of latitude 50º N), eastern China and the eastern Himalayas.

Irano-Turanian Region (IT): From central Anatolia, north-eastern Syria and the Caspian seashores in the East (including south-eastern Transcaucasia) throughout northern Iraq and the Iranian Plateau to the Gobi Desert in the East. It is bordered on the North by WP and EP.

Indian-Indochinese Region (IN): It comprises the Indian Plate and the Indochinese Peninsula.

African Region (AF): It extends throughout the Sahara Desert, from the Atlantic Ocean to Egypt, southern Palestine, Syrian Desert, southern Iraq, the Arabian Peninsula, shores of the Persian Gulf, the tropical deserts of southern Iran, Pakistan and north-western India. It also includes the rest of continental Africa North of the equator, except tropical rainforests.

Macaronesian Region (MC): The archipelagos of Azores, Madeira, Savages, Canary Islands and Cape Verde.

Mediterranean Region (ME): The entire Mediterranean Basin, including all Mediterranean islands and coastal areas. In Europe, it includes the Iberian Peninsula (except northern regions included in WP), south-eastern France and the South of the Italian and Balkan Peninsulas. In Africa, it extends from Morocco through northern Algeria and Tunisia to northern Libya (including coastal Cyrenaica). It also extends to the coasts of northern Palestine, Lebanon, western Syria and western Anatolia in the East.

Nearctic Region (NE): It covers most of North America (excluding the Madrean Region), from the Atlantic coast in the East to east-central Texas, north-eastern New Mexico, most of the Rocky Mountains and Sierra Nevada (California) in the West.

Madrean Region (MA): It covers south-western North America, from the south-western Rocky Mountains in the Northeast to the Pacific coasts of California and Mexico in the West, and the Mexican Sierra Madre in the South (it excludes the Balsas River basin and Mexican southern lowlands).

Neotropical Region (NT): It includes the Gulf of Mexico lowlands (South of Tampico), the southern tip of Florida, the western coastline of Venezuela, the Revillagigedo and Galapagos archipelagos in the Pacific Ocean as well as the South American western coast from Mexico (around Sinaloa) to Tierra del Fuego, the Andean Mountains and Patagonia.

*Time stratified approach*

Following Meseguer, Lobo, Ree, Beerling, and Sanmartín (2015), four time slices were established in order to incorporate paleogeographic changes in biogeographic reconstructions by assigning specific dispersal probability values between areas for each time interval.

The first time slice (TSI), from 60 to 35 Ma, is a period characterized by the proximity of all northern landmasses (Meseguer et al., 2015) allowing the presence of a continuous "boreotropical" flora due to stable temperatures and warm climates at high latitudes during the Early Eocene (Wolfe, 1975). The second time slice (TSII), covering 35 to 10 Ma, is defined by low temperatures resulting in the development of a mixed-mesophytic forest and the retraction of some of the tropical flora to low latitudes mainly in Asia (Meseguer et al., 2015; Morley, 2007), the closure of the Turgai strait allowing land connection between WP and EP (Sanmartín, Enghoff, & Ronquist, 2001) and the collision between the Caribbean and North American plates starting to uplift Central American lands (Morley, 2003). During the third time slice (TSIII), from 10 to 3.5 Ma, the uplift of a land bridge between North America and South America connected these two areas (Morley, 2003) and the collision of the Indian Plate against the Eurasian and Arabian plates completed the uplift of the Irano-Turanian Region (Manafzadeh, Salvo, & Conti, 2014). The fourth time slice (TSIV), spanning 3.5 to 0 Ma, started with the opening of the Bering strait causing the breakup of the land connection between the Eastern Palearctic and Nearctic regions (Meseguer et al., 2015; Sanmartín et al., 2001), and is also characterised by the glacial and inter-glacial cycles of the Pleistocene (Médail & Diadema, 2009). Dispersal to Macaronesia was considered in all time slices accounting for the times of origin estimated by Fernández-Palacios et al. (2011) for each archipelago.

Dispersal probability values between pairs of areas were established for each time slice and scaled to represent connection between areas. Dispersal was not limited (scalar = 1) when areas were directly connected by land; it was set as less probable (scalar = 0.5) for adjacent areas not directly connected by land; even less likely dispersal (scalar = 0.1) was applied to non-adjacent and very distant areas; and dispersal was set to null (scalar = 0) when any of the two landmasses was below sea level. Fig. S1 shows the matrices with the probability values for each time slice.

### RESULTS

#### *Description of major clades within the Antirrhineae*

The main clades within the tribe were well defined and congruent with previous phylogenies (Fernández-Mazuecos et al., 2019). *Anarrhinum* (B), *Kickxia* (C) and

*Nanorrhinum* (D) were estimated to form an early diverging clade. Sister to this, a group with amphiatlantic distribution formed by the New World *Maurandya* clade (*Epixiphium*, *Holmgrenanthe*, *Maurandya*, *Maurandella*, *Rhodochiton*, *Lophospermum* and *Mabrya*) (E) plus the Old World *Cymbalaria* – *Asarina* – *Gadoria* clade (F) diverges from another group containing the rest of taxa. Branching off these remaining taxa is another Old World/New World lineage represented by *Schweinfurthia* (G) plus the *Galvezia* – *Pseudorontium* clade (H), leaving out a large group of taxa containing five main lineages. The *Chaenorhinum* clade (*Chaenorhinum* and *Albraunia*) (I) is the first lineage to diverge within this large group, and the remaining sister group splits in two descendants, one leading to the ancestor of *Antirrhinum* (J), the *Pseudomisopates* – *Acanthorrhinum* – *Misopates* clade (K), and the *Sairocarpus* clade (*Sairocarpus*, *Mohavea*, *Howelliella*, *Neogaerrhinum* and *Gambelia*) (L), and another one leading to *Linaria* (M), the most widespread and diverse of the Antirrhineae genera.

**Table S1.** GenBank accession numbers for previously published and newly generated DNA sequences of Antirrhineae and the outgroup used in the present study.

| **Taxon** | **ITS** | ***rpl*32-*trn*L** | ***ndh*F** |
|---|---|---|---|
| **OUTGROUP** | | | |
| ACANTHACEAE | | | |
| *Thunbergia alata* Bojer ex Sims | - | - | U12667 |
| BIGNONIACEAE | | | |
| *Catalpa* Scop. | - | - | L36397 |
| *Kigelia africana* (Lam.) Benth. | - | - | AF102632 |
| CALCEOLARIACEAE | | | |
| *Calceolaria* L. | - | - | AF123679 |
| GESNERIACEAE | | | |
| *Nematanthus hirsutus* (Mart.) Wiehler | - | - | L36404 |
| *Streptocarpus holstii* Engl. | - | - | L36415 |
| LAMIACEAE | | | |
| *Lamium purpureum* L. | - | - | U78694 |
| *Teucrium fruticans* L. | - | - | U78686 |
| OLEACEAE | | | |
| *Fraxinus chinensis* Roxb. | - | - | DQ673275 |
| *Jasminum mesnyi* Hance | - | - | DQ673267 |
| *Ligustrum vulgare* L. | - | - | AF130164 |
| *Olea europaea* L. | - | - | DQ673278 |
| OROBANCHACEAE | | | |
| *Bartsia alpina* L. | - | - | AF123678 |
| *Pedicularis foliosa* L. | - | - | AF123689 |
| PEDALIACEAE | | | |
| *Sesamum indicum* L. | - | - | L36413 |
| PLANTAGINACEAE | | | |
| *Bacopa caroliniana* (Walter) B.L. Rob. | - | - | AF123677 |
| *Callitriche hermaphroditica* L. | - | - | L36396 |
| *Chelone obliqua* L. | - | - | AF123680 |
| *Collinsia grandiflora* Douglas ex Lindl. | HQ653045 | - | AF188182 |
| *Digitalis grandiflora* Mill. | - | - | L36399 |
| *Digitalis purpurea* L. | - | - | AF130150 |
| *Globularia nudicaulis* L. | - | - | AF123681 |
| *Gratiola pilosa* Michx. | - | - | AF188183 |
| *Isoplexis canariensis* (L.) Loudon | - | - | AJ617597 |
| *Lafuentea jeanpertiana* Maire | MK095630 | MK095683 | MK095671 |
| *Lafuentea rotundifolia* Lag. | AF509816 | MK095684 | JN848475 |
| *Penstemon gracilis* Nutt. | DQ531102 | - | KT176851 |
| *Plantago lanceolata* L. | - | - | AF130151 |
| *Stemodia suffruticosa* Kunth | - | - | EF527455 |
| *Veronica persica* Poir. | - | - | L36419 |
| PLOCOSPERMATACEAE | | | |
| *Plocosperma buxifolium* Benth. | - | - | AJ011985 |
| SCROPHULARIACEAE | | | |
| *Buddleja davidii* Franch | - | - | AF130143 |
| *Scrophularia* L. | - | - | L36411 |
| VERBENACEAE | | | |

| **Taxon** | **ITS** | ***rpl*32-*trn*L** | ***ndh*F** |
|---|---|---|---|
| *Verbena bracteata* Lag. & Rodr. | - | - | L36418 |
| **INGROUP (PLANTAGINACEAE, ANTIRRHINEAE)** | | | |
| *Acanthorrhinum ramosissimum* (Coss. & Durieu) Rothm. | AY731261 | KM104746 | JN848476 |
| *Albraunia foveopilosa* Speta | AY731250 | KT031976 | JN848477 |
| *Anarrhinum bellidifolium* (L.) Wild. | AY731263 | KP851136 | JN848478 |
| *Anarrhinum corsicum* Jord. & Fourr | - | **forthcoming** | AJ245815 |
| *Anarrhinum duriminium* (Brot.) Pers. | MK095631 | MK095685 | - |
| *Anarrhinum forskaohlii* (J.F.Gmel.) Cufod | MK095632 | MK095686 | - |
| *Anarrhinum fruticosum* Desf. | AF513881 | - | - |
| *Anarrhinum laxiflorum* Boiss. | KT187718 | MK095687 | KT187751 |
| *Anarrhinum longipedicellatum* R.Fernandes | - | MK095688 | - |
| *Anarrhinum pedatum* Desf. | MK095633 | MK095689 | - |
| *Antirrhinum australe* Rothm. | AY731273 | **forthcoming** | KT187752 |
| *Antirrhinum braun-blanquetii* Rothm. | AY731269 | **forthcoming** | KT187753 |
| *Antirrhinum charidemi* Lange | AY731282 | **forthcoming** | KT187754 |
| *Antirrhinum cirrhigerum* (Welw. ex Ficalho) Rothm. | EU677200 | **forthcoming** | KT187755 |
| *Antirrhinum controversum* Pau | AY731272 | **forthcoming** | KT187756 |
| *Antirrhinum graniticum* Rothm. | AY731283 | JF694120 | KT187757 |
| *Antirrhinum grosii* Font Quer | AY731281 | **forthcoming** | KT187758 |
| *Antirrhinum hispanicum* Chav. | AY731286 | **forthcoming** | KT187759 |
| *Antirrhinum latifolium* Mill. | AY731274 | **forthcoming** | KT187760 |
| *Antirrhinum linkianum* Boiss. & Reut. | AY731278 | **forthcoming** | KT187773 |
| *Antirrhinum litigiosum* Pau | AY731271 | **forthcoming** | KT187761 |
| *Antirrhinum lopesianum* Rothm. | EU677217 | **forthcoming** | KT187762 |
| *Antirrhinum majus* L. | AY731280 | KP851132 | JN848479 |
| *Antirrhinum meonanthum* Hoffmans. & Link | AY731284 | **forthcoming** | KT187763 |
| *Antirrhinum microphyllum* Rothm. | AY731267 | **forthcoming** | KT187764 |
| *Antirrhinum molle* L. | AY731268 | **forthcoming** | KT187765 |
| *Antirrhinum mollissimum* (Pau) Rothm. | AY731275 | **forthcoming** | KT187766 |
| *Antirrhinum onubense* (Fern.Casas) Fern.Casas | MK095634 | **forthcoming** | MK095672 |
| *Antirrhinum pertegasii* Pau ex Rothm. | EU677226 | **forthcoming** | KT187767 |
| *Antirrhinum pulverulentum* Lázaro Ibiza | AY731279 | **forthcoming** | KT187768 |
| *Antirrhinum rothmaleri* (P.Silva) Amich, Bernardos & García-Barriuso | MK095635 | **forthcoming** | MK095673 |
| *Antirrhinum sempervirens* Lapeyr. | AY731270 | **forthcoming** | KT187769 |
| *Antirrhinum siculum* Mill. | AY731276 | **forthcoming** | KT187770 |
| *Antirrhinum subbaeticum* Güemes, Mateu & Sánchez-Gómez | AY731287 | **forthcoming** | MK095674 |
| *Antirrhinum tortuosum* Bosc ex Vent. | AY731285 | **forthcoming** | KT187771 |
| *Antirrhinum valentinum* Font Quer | AY731266 | **forthcoming** | KT187772 |
| *Asarina procumbens* Mill. | AF513879 | KP851129 | KT187774 |
| *Chaenorhinum calycinum* (Banks & Sol.) P.H.Davis | MK095636 | MK095690 | - |
| *Chaenorhinum crassifolium* (Cav.) Kostel. | KT187719 | KP851135 | KT187775 |
| *Chaenorhinum cryptarum* (Boiss. & Hausskn.) P.H.Davis | KT187720 | - | KT187776 |
| *Chaenorhinum exile* (Coss. & Kralik) Lange | MK095637 | MK095691 | KT187786 |
| *Chaenorhinum flexuosum* (Desf.) Lange | MK095638 | MK095692 | - |
| *Chaenorhinum formenterae* Gand. | MK095639 | MK095693 | - |
| *Chaenorhinum glareosum* (Boiss.) Willk | KT187721 | KT031951 | KT187777 |
| *Chaenorhinum grandiflorum* (Coss.) Willk | KT187722 | KT031947 | KT187778 |
| *Chaenorhinum huber-morathii* P.H.Davis | MK095640 | MK095694 | - |
| *Chaenorhinum johnstonii* (Stapf) Pennell | KT031890 | KT031948 | - |
| *Chaenorhinum litorale* (Willd.) Rouy | MK095641 | MK095695 | - |

| **Taxon** | **ITS** | ***rpl*32-*trn*L** | ***ndh*F** |
|---|---|---|---|
| *Chaenorhinum litorale* subsp. *pterosporum* (Fisch. & C.A.Mey.) P.H.Davis | KT031883 | MK095696 | - |
| *Chaenorhinum macropodum* Lange | KT187723 | JF694119 | KT187779 |
| *Chaenorhinum minus* (L.) Lange | KT187724 | KT031946 | KT187780 |
| *Chaenorhinum origanifolium* (L.) Kostel | KT187725 | KT031941 | KT187781 |
| *Chaenorhinum raveyi* (Boiss.) Pau | KT187729 | KT031943 | KT187785 |
| *Chaenorhinum reyesii* (C.Vicioso & Pau) Benedí | MK095642 | MK095697 | MK095675 |
| *Chaenorhinum robustum* Loscos | KT187727 | KT031942 | KT187783 |
| *Chaenorhinum rubrifolium* (Robill. & Castagne ex DC.) Fourr. | KT187728 | KT031944 | KT187784 |
| *Chaenorhinum segoviense* (Reut. ex Rouy) Rouy | KT187726 | MK095698 | KT187782 |
| *Chaenorhinum serpyllifolium* (Lange) Lange | KT187731 | MK095699 | KT187787 |
| *Chaenorhinum spicatum* Korovin | AY731258 | KT031949 | JN848483 |
| *Chaenorhinum tenellum* (Cav.) Lange | KT187732 | MK095700 | KT187788 |
| *Chaenorhinum tuberculatum* Speta | KT031887 | KT031945 | - |
| *Chaenorhinum villosum* (L.) Lange | KT187733 | KT031940 | KT187789 |
| *Cymbalaria aequitriloba* (Viv.) A.Chev. | KP735225 | KP851100 | KT187792 |
| *Cymbalaria fragilis* (Rodrig.) A.Chev | KP735211 | KP851090 | KP851004 |
| *Cymbalaria glutinosa* Bigazzi & Rafaelli | KP735216 | KP851114 | KP851029 |
| *Cymbalaria glutinosa* subsp. *brevicalcarata* Bigazzi & Rafaelli | KP735218 | KP851105 | - |
| *Cymbalaria hepaticifolia* Wettst. | KP735215 | KP851099 | KP851013 |
| *Cymbalaria longipes* (Boiss. & Heldr.) A.Chev. | KP735232 | KP851123 | KP851038 |
| *Cymbalaria microcalyx* (Boiss.) Wettst. | KP735238 | - | KP851041 |
| *Cymbalaria microcalyx* subsp. *acutiloba* (Boiss. & Heldr.) W.Greuter | KP735212 | KP851126 | - |
| *Cymbalaria microcalyx* subsp. *dodekanesii* Greuter | KP735208 | KP851127 | - |
| *Cymbalaria microcalyx* subsp. *ebelii* (Cufod.) Cufod. | KP735236 | KP851121 | - |
| *Cymbalaria microcalyx* subsp. *minor* (Cufod.) Greuter | KP735237 | KP851122 | - |
| *Cymbalaria muelleri* (Moris.) A.Chev | KP735210 | KP851098 | KP851012 |
| *Cymbalaria muralis* G.Gaertn., B.Mey. & Scherb. | KP735231 | KP851102 | KT187790 |
| *Cymbalaria muralis* subsp. *visianii* (Jáv.) D.A.Webb | KP735227 | KP851116 | - |
| *Cymbalaria pallida* Wettst. | KP735235 | KP851119 | KT187791 |
| *Cymbalaria pubescens* (J.Presl & C.Presl) Cufod. | KP735229 | KP851106 | KP851021 |
| *Epixiphium wislizeni* (Engelm. ex A.Gray) Munz | AY878930 | KP851130 | KP851046 |
| *Gadoria falukei* Güemes & Mota | MK095643 | MK095701 | - |
| *Galvezia ballii* Munz | KT187736 | **forthcoming** | KT187793 |
| *Galvezia elisensii* M.O.Dillon & Quip. | KX257937 | **forthcoming** | KX257890 |
| *Galvezia fruticosa* J.F.Gmel. | KT187737 | KP851128 | KX257911 |
| *Galvezia lanceolata* Pennell | KT187738 | **forthcoming** | KT187794 |
| *Galvezia leucantha* Wiggins | KX257944 | **forthcoming** | KX257921 |
| *Galvezia leucantha* subsp. *porphyrantha* A.Tye & H.Jäger | KX257950 | - | KX257923 |
| *Galvezia leucantha* subsp. *pubescens* Wiggins | KX257990 | - | KX257925 |
| *Gambelia juncea* (Beneth.) D.A.Sutton | AY316310 | - | KT187795 |
| *Gambelia speciosa* Nutt. | KT187739 | - | KT187796 |
| *Holmgrenanthe petrophila* (Coville & C.V.Morton) Elisens | AY880231 | - | - |
| *Howelliella ovata* (Eastw.) Rothm. | AF513899 | **forthcoming** | AJ250385 |
| *Kickxia aegyptiaca* (L.) Nábelek | KT187741 | KT031964 | KT187799 |
| *Kickxia aegyptiaca* subsp. *fruticosa* (Desf.) Wickens | **forthcoming** | **forthcoming** | - |
| *Kickxia caucásica* (Muss.Puschk. ex Spreng.) Kuprian. | KT031895 | KT031953 | - |
| *Kickxia cirrhosa* (L.) Fritsch | MK095644 | KT031954 | - |
| *Kickxia commutata* (Bernh. ex Rchb.) Fritsch | KT031897 | KT031955 | - |
| *Kickxia elatine* (L.) Dumort. | AY731265 | KT031956 | AJ245816 |
| *Kickxia elatine* subsp. *crinita* (Mabille) Greuter | KT031899 | KT031957 | - |

| **Taxon** | **ITS** | ***rpl*32-*trn*L** | ***ndh*F** |
|---|---|---|---|
| *Kickxia floribunda* (Boiss.) Täckh. & Boulos | KT031900 | KT031958 | - |
| *Kickxia lanigera* (Desf.) Hand.-Mazz. | KT187740 | KT031965 | KT187797 |
| *Kickxia scariosa* (Lam.) Rothm | - | KT031969 | - |
| *Kickxia sieberi* (Rchb.) Dörfl & Allan | KT031912 | KT031971 | - |
| *Kickxia spuria* (L.) Dumort. | AY731264 | KT031973 | KT187798 |
| *Kickxia spuria* subsp. *integrifolia* (Brot.) R.Fern | KT031913 | KT031972 | - |
| *Linaria accitensis* L.Sáez, Juan, M.B.Crespo, F.B.Navarro, J.Peñas & Roquet | MK095645 | MK095702 | - |
| *Linaria aeruginea* (Gouan) Cav. | JQ814486 | JN663397 | KT187836 |
| *Linaria albifrons* (Sibth. & Sm.) Steudel | JX481129 | JQ814606 | KT187808 |
| *Linaria algarviana* Chav. | JX481086 | JF694130 | KT187800 |
| *Linaria almijarensis* Campo & Amo | KF623175 | JN663410 | - |
| *Linaria alpina* (L.) Mill. | JQ814489 | JN663403 | KT187809 |
| *Linaria altaica* Fisch. ex Kuprian. | MK095646 | MK095703 | - |
| *Linaria amethystea* (Vent.) Hoffmanns. & Link | JQ814490 | JN663405 | AJ250386 |
| *Linaria amethystea* subsp. *broussonetii* (Poir.) Malato-Beliz | KF623180 | JN663406 | - |
| *Linaria amoi* Campo ex Amo | JX481108 | JN663407 | KT187801 |
| *Linaria angustissima* (Loisel.) Borbás | JX481133 | MK095704 | - |
| *Linaria anticaria* Boiss. & Reuter | JQ814491 | JN663408 | - |
| *Linaria antilibanotica* Rech.f. | JX481134 | MK095705 | - |
| *Linaria arenaria* DC. | JX481112 | JN663414 | - |
| *Linaria armeniaca* Chav. | JX481080 | MK095706 | - |
| *Linaria arvensis* (L.) Desf. | JQ814494 | JN663415 | - |
| *Linaria atlantica* Boiss. & Reut. | - | JN663419 | - |
| *Linaria atrofusca* Rouy | KF623182 | JN663392 | - |
| *Linaria azerbaijanensis* Hamdi & Assadi | KJ747000 | - | - |
| *Linaria badalii* Loscos | JQ814495 | JN663421 | - |
| *Linaria baligaliensis* Patzak | JX481142 | MK095707 | - |
| *Linaria bamianica* Patzak | KJ747001 | MK095708 | - |
| *Linaria becerrae* Blanca, Cueto & J.Fuentes | JX481087 | JF694170 | KT187830 |
| *Linaria benitoi* Fern.Casas | - | JN663491 | - |
| *Linaria bipartita* Fern.Casas | JX481094 | JF694131 | KT187811 |
| *Linaria bipunctata* (L.) Chaz. | JQ814496 | JN663423 | - |
| *Linaria bordiana* Santa & Simonneau | KC994568 | KC994580 | - |
| *Linaria bordiana* subsp. *kralikiana* (Maire) D.A.Sutton | KC994569 | JF694134 | - |
| *Linaria boushehrensis* Hamdi & Assadi | KJ747025 | - | - |
| *Linaria brachyceras* (Bunge) Kuprian. | **forthcoming** | **forthcoming** | - |
| *Linaria bubanii* Font Quer | JQ814537/8 | JN663424 | - |
| *Linaria caesia* (Pers.) F.Dietr. | JX481105 | JN663425 | - |
| *Linaria canadensis* (L.) Dum.Cours. | AY883085 | KT031924 | - |
| *Linaria capraria* Moris & De Not. | JX481146 | KT031925 | - |
| *Linaria cavanillesii* Chav. | KP735198 | KP851134 | KP851050 |
| *Linaria chalepensis* (L.) Mill. | JX481081 | JF694127 | KT187812 |
| *Linaria clementei* Haens. | JX481089 | JF694135 | KT187813 |
| *Linaria confertiflora* Benth. | MK095647 | MK095709 | - |
| *Linaria corifolia* Desf. | JX481135 | MK095710 | KT187815 |
| *Linaria cretacea* Fisch. ex Spreng. | KT031851 | KT031919 | - |
| *Linaria creticola* Kuprian. | JX481143 | MK095711 | - |
| *Linaria dalmatica* (L.) Mill. | JX481136 | JQ814607 | KT187816 |
| *Linaria damascena* Boiss. & Gaill. | MK095648 | MK095712 | - |
| *Linaria decipiens* Batt. | JX481125 | MK095713 | - |

| **Taxon** | **ITS** | ***rpl*32-*trn*L** | ***ndh*F** |
|---|---|---|---|
| *Linaria depauperata* Leresche ex Lange | JX481107 | JN663431 | KT187802 |
| *Linaria depauperata* subsp. *hegelmaieri* (Lange) De la Torre, Alcaraz & M.B.Crespo | KC625799 | - | - |
| *Linaria depauperata* subsp. *ilergabona* (M.B.Crespo & Arán) L.Sáez | KF623189 | - | - |
| *Linaria diffusa* Hoffmanns. & Link | JX481114 | JN663432 | - |
| *Linaria elegans* Cav. | JX481103 | KC521097 | KT187817 |
| *Linaria elymaitica* (Boiss.) Kuprian. | KJ747002 | - | - |
| *Linaria farsensis* Hamdi & Assadi | KJ747003 | - | - |
| *Linaria fastigiata* Chav. | KJ747004 | MK095714 | - |
| *Linaria faucicola* Leresche & Levier | JX481117 | JN663436 | - |
| *Linaria ficalhoana* Rouy | MK095649 | MK095715 | - |
| *Linaria filicaulis* Boiss. ex Leresche & Levier | JQ814500 | JN663434 | - |
| *Linaria flava* (Poiret) Desf. | JX481130 | JQ814608 | KT187818 |
| *Linaria galioides* Ball | KC994575 | JF694153 | - |
| *Linaria gattefossei* Maire & Weiller | KC994571 | KC994581 | - |
| *Linaria genistifolia* (L.) Mill. | JX481137 | JQ814609 | KT187834 |
| *Linaria gharbensis* Batt. & Pit. | JX481100 | JF694139 | KT187819 |
| *Linaria glacialis* Boiss. | JQ814505 | KC625787 | MK095676 |
| *Linaria glauca* (L.) Chaz. | JX481116 | JN663439 | - |
| *Linaria glauca* subsp. *olcadium* Valdés & D.A.Webb | JQ814506 | JN663441 | - |
| *Linaria golestanensis* Hamdi & Assadi | KJ747029 | - | - |
| *Linaria grandiflora* Desf. | JX481151 | MK095716 | KT187820 |
| *Linaria griffithsii* Benth. | MK095650 | MK095717 | - |
| *Linaria guilanensis* Hamdi & Assadi | KJ747005 | - | - |
| *Linaria haelava* (Forssk.) F.G.Dietr. | JX481122 | JQ814610 | - |
| *Linaria hellenica* Turrill | JX481102 | JF694140 | - |
| *Linaria hirta* (Loefl. ex L.) Moench | JX481120 | MK095718 | KT187822 |
| *Linaria huteri* Lange | JX481111 | JN663442 | - |
| *Linaria iconia* Boiss. & Heldr. | JX481154 | MK095719 | - |
| *Linaria imzica* Gómiz | KC878736 | JF694141 | - |
| *Linaria incarnata* (Vent.) Spreng. | JX481088 | JF694145 | KT187803 |
| *Linaria incompleta* Kuprian. | MK095651 | MK095720 | - |
| *Linaria intricata* Coincy | JX481113 | JN663427 | - |
| *Linaria iranica* Hamdi & Assadi | KJ747031 | - | - |
| *Linaria japonica* Miq. | JX481141 | MK095721 | - |
| *Linaria joppensis* Bornm. | JX481123 | JQ814611 | - |
| *Linaria karajensis* Hamdi & Assadi | KJ747006 | - | - |
| *Linaria kavirensis* Hamdi & Assadi | KJ747032 | - | - |
| *Linaria khalkhalensis* Hamdi & Assadi | KJ747007 | - | - |
| *Linaria khorasanensis* Hamdi & Assadi | KJ747008 | - | - |
| *Linaria kurdica* Boiss. & Hohen. | JX481139 | MK095722 | - |
| *Linaria kurdica* subsp. *pycnophylla* (Boiss. & Bal.) P.H.Davis | KJ747010 | KT026488 | - |
| *Linaria latifolia* Desf. | JX481124 | MK095723 | - |
| *Linaria laxiflora* Desf. | JX481128 | JQ814612 | - |
| *Linaria leptoceras* Kuprian. | KJ747011 | - | - |
| *Linaria lilacina* Lange | JX481156 | JN663444 | KT187835 |
| *Linaria lineolata* Boiss. | KJ747012 | MK095724 | - |
| *Linaria loeselii* Schweigg. | JX481153 | JQ814613 | - |
| *Linaria macroura* (M.Bieb.) M.Bieb. | MK095652 | MK095725 | - |
| *Linaria mamorensis* Mazuecos, Vigalondo & L.Sáez | KC878745 | JF694142 | - |
| *Linaria maroccana* Hook.f. | JX481097 | JF694149 | KT187823 |

| **Taxon** | **ITS** | ***rpl*32-*trn*L** | ***ndh*F** |
|---|---|---|---|
| *Linaria mazandaranensis* Hamdi & Assadi | KJ747033 | - | - |
| *Linaria meyeri* Kuprian. | JX481150 | Q814614 | KT187824 |
| *Linaria michauxii* Chav. | KJ747013 | - | - |
| *Linaria micrantha* (Cav.) Hoffmanns. & Link | JQ814513 | JN663445 | KT187825 |
| *Linaria microsepala* A.Kern. | MK095653 | MK095726 | - |
| *Linaria multicaulis* (L.) Mill. | JX481098 | JF694150 | - |
| *Linaria multicaulis* subsp. *aurasiaca* (Pomel) D.A.Sutton | KC994573 | JF694151 | - |
| *Linaria multicaulis* subsp. *heterophylla* (Desf.) D.A.Sutton | KC994576 | JF694157 | KC878830 |
| *Linaria munbyana* Boiss. & Reut. | JQ814515 | JN663447 | - |
| *Linaria nevadensis* (Boiss.) Boiss. & Reut. | JQ814487 | JN663401 | - |
| *Linaria nigricans* Lange | JX481104 | JF694186 | KT187826 |
| *Linaria nivea* Boiss. & Reut. | JX481155 | KF623211 | MK095677 |
| *Linaria nurensis* Boiss. & Hausskn. ex Boiss. | MK095654 | MK095727 | - |
| *Linaria oblongifolia* (Boiss.) Boiss. & Reut. | JQ814516 | JN663449 | - |
| *Linaria oblongifolia* subsp. *haenseleri* (Boiss. & Reuter) Valdès | KF623197 | JN663448 | - |
| *Linaria odora* (M.Bieb.) Fisch. | JX481152 | JQ814615 | - |
| *Linaria oligantha* Lange | JX481121 | JN663450 | - |
| *Linaria onubensis* Pau | KC878751 | KC878769 | - |
| *Linaria orbensis* Carretero & Boira | JQ814518 | JN663453 | - |
| *Linaria orientalis* Hamdi & Assadi | KJ747035 | - | - |
| *Linaria parviracemosa* D.A.Sutton | - | MK095728 | - |
| *Linaria pedunculata* (L.) Chaz. | JX481101 | JF694161 | - |
| *Linaria pelisseriana* (L.) Mill. | JX481082 | MK095729 | KT187828 |
| *Linaria peloponnesiaca* Boiss. & Heldr. | JX481148 | JQ814616 | KT187821 |
| *Linaria peltieri* Batt. | - | MK095730 | - |
| *Linaria pinifolia* (Poir.) Thell. | KC878753 | JF694168 | - |
| *Linaria platycalyx* Boiss. | JQ814520 | JN663454 | KT187804 |
| *Linaria polygalifolia* Hoffmanns. & Link | JQ814522 | JN663456 | KT187805 |
| *Linaria polygalifolia* subsp. *aguillonensis* (García Mart.) S.Castroviejo & E.Lago | KF623200 | JN663455 | - |
| *Linaria polygalifolia* subsp. *lamarckii* (Rouy) D.A.Sutton | JQ814522 | JQ814617 | - |
| *Linaria popovii* Kuprian. | MK095655 | MK095731 | - |
| *Linaria propinqua* Boiss. & Reut | JQ814524 | JN663460 | - |
| *Linaria pseudoviscosa* Murb. | JX481099 | JF694169 | - |
| *Linaria purpurea* (L.) Mill. | **forthcoming** | **forthcoming** | - |
| *Linaria pyramidalis* (Vent.) F .G.Dietr. | KJ747017 | MK095732 | - |
| *Linaria pyramidalis* subsp. *kopetdaghensis* (Kuprian.) D.A.Sutton | KJ747016 | KT026497 | - |
| *Linaria reflexa* (L.) Chaz. | JX481126 | KF623209 | KT187829 |
| *Linaria remotiflora* Patzak | KJ747018 | - | - |
| *Linaria repens* (L.) Mill. | JX481144 | KF623210 | - |
| *Linaria ricardoi* Cout. | MK095656 | MK095733 | - |
| *Linaria rubioides* Vis. & Pančić | JX481149 | MK095734 | - |
| *Linaria salzmannii* Boiss. | KC994579 | KC994582 | - |
| *Linaria saturejoides* Boiss. | JQ814525 | JN663462 | - |
| *Linaria saxatilis* (L.) Chaz. | JX481115 | JN663463 | - |
| *Linaria schelkownikowii* Schischk. | - | MK095735 | - |
| *Linaria schirvanica* Fomin | MK095657 | MK095736 | - |
| *Linaria sessilis* Kuprian. | **forthcoming** | **forthcoming** | - |
| *Linaria shahroudensis* Hamdi & Assadi | KJ747019 | - | - |
| *Linaria simplex* Willd. ex Desf. | JQ814528 | JN663474 | - |
| *Linaria spartea* (L.) Chaz. | JX481090 | JF694171 | KT187831 |

| Taxon | ITS | *rpl*32-*trn*L | *ndh*F |
|---|---|---|---|
| *Linaria striatella* Kuprian. | MK095658 | MK095737 | |
| *Linaria subandina* Diels | JX481084 | MK095738 | |
| *Linaria supina* (L.) Chaz. | JQ814530 | JN663481 | |
| *Linaria supina* subsp. *maritima* (DC.) M.Lainz | KF623203 | - | - |
| *Linaria tenuis* (Viv.) Spreng. | JX481096 | JF694174 | |
| *Linaria texana* Scheele | JX481085 | JN663496 | JN848491 |
| *Linaria thibetica* Franch. | JX481140 | JQ814618 | - |
| *Linaria thymifolia* (Vahl) OC. | JX481106 | JN663482 | - |
| *Linaria tingitana* Boiss. & Reut. | JX481092 | JF694175 | - |
| *Linaria transiliensis* Kuprian. | MK095659 | MK095739 | - |
| *Linaria triornithophora* (L.) Cav. | JX481083 | JF694128 | KT187832 |
| *Linaria triphylla* (L.) Mill. | JX481132 | JQ814619 | - |
| *Linaria tristis* (L.) Mill. | JX481109 | JN663490 | - |
| *Linaria tristis* subsp. *lurida* (Ball) Maire | - | JN663484 | - |
| *Linaria tristis* subsp. *mesatlantica* D.A.Sutton | KF623205 | JN663486 | - |
| *Linaria tristis* subsp. *pectinata* (Pau & Font Quer) Maire | KF623206 | JN663488 | - |
| *Linaria tursica* Valdés & Cabezudo | JQ814533 | JN663492 | KT187833 |
| *Linaria unaiensis* Patzak | MK095660 | MK095740 | - |
| *Linaria venosa* Lindl. | - | MK095741 | - |
| *Linaria ventricosa* Coss. & Balansa | JX481145 | JQ814620 | KT187814 |
| *Linaria verticillata* Boiss. | JX481110 | JN663493 | KT187806 |
| *Linaria verticillata* subsp. *cuartanensis* (Degen & Hervier) L.Sáez & M.B.Crespo | KF623188 | JN663409 | - |
| *Linaria virgata* (Poir.) Desf. | JX481131 | MK095742 | - |
| *Linaria viscosa* (L.) Chaz. | JX481091 | JF694178 | KT187807 |
| *Linaria vulgaris* Mill. | JX481138 | JQ814621 | JN848484 |
| *Linaria warionis* Pomel | JX481127 | JQ814622 | - |
| *Linaria weilleri* Emb. & Maire | JX481093 | JF694182 | - |
| *Lophospermum breedlovei* Elisens | MK095661 | **forthcoming** | MK095678 |
| *Lophospermum erectum* (Hemsl.) Rothm. | KT187742 | **forthcoming** | - |
| *Lophospermum erubescens* D.Don | AY731249 | **forthcoming** | JN848485 |
| *Lophospermum purpusii* (Brandegee) Rothm. | KC782760 | **forthcoming** | - |
| *Lophospermum scandens* D.Don | MK095662 | **forthcoming** | - |
| *Lophospermum turneri* Elisens | MK095663 | **forthcoming** | MK095679 |
| *Mabrya acerifolia* (Pennell) Elisens | KT187743 | **forthcoming** | KT187837 |
| *Mabrya coccinea* (I.M.Johnst.) Elisens | MK095664 | **forthcoming** | MK095680 |
| *Mabrya geniculata* (B.L.Rob. & Fernald) Elisens | MK095665 | - | - |
| *Mabrya rosei* (Munz) Elisens | KT187744 | **forthcoming** | KT187838 |
| *Maurandella antirrhiniflora* (Willd.) Rothm. | KT187745 | **forthcoming** | JN848487 |
| *Maurandya barclaiana* Lindl. | KT187746 | **forthcoming** | KT187839 |
| *Maurandya scandens* (Cav.) Pers. | JX481079 | **forthcoming** | KT187840 |
| *Misopates calycinum* (Vent.) Rothm. | AY731259 | **forthcoming** | KT187842 |
| *Misopates chrysothales* (Font Quer) Rothm. | KT187747 | - | KT187841 |
| *Misopates microcarpum* (Pomel) D.A.Sutton | MK095666 | **forthcoming** | MK095681 |
| *Misopates orontium* (L.) Raf. | AY731260 | KP851133 | JN848488 |
| *Misopates salvagense* D.A.Sutton | MK095667 | - | MK095682 |
| *Mohavea breviflora* Coville | AF513892 | - | KT187843 |
| *Mohavea confertiflora* (Benth.) A.Heller | AF513891 | KP851138 | AJ250389 |
| *Nanorrhinum acerbianum* (Boiss.) Betsche | KT031894 | KT031952 | - |
| *Nanorrhinum asparagoides* (Schweinf.) Ghebr. | MK095668 | MK095743 | - |
| *Nanorrhinum cabulicum* (Benth.) Podlech & Iranshahr | KT031916 | KT031975 | - |

| **Taxon** | **ITS** | ***rpl*32-*trn*L** | ***ndh*F** |
|---|---|---|---|
| *Nanorrhinum elegans* (G.Forst.) Ghebr. | KT187748 | MK095744 | JN848489 |
| *Nanorrhinum heterophyllum* (Schousb.) Ghebr. | MK095669 | MK095745 | KT187844 |
| *Nanorrhinum heterophyllum* subsp. *canariensis* (V. W. Smith) | - | KT031960 | - |
| *Nanorrhinum judaicum* (Danin) Yousefi & Zarre | KT031907 | KT031966 | - |
| *Nanorrhinum macilentum* (Decne.) Betsche | KT031908 | KT031967 | - |
| *Nanorrhinum ovatum* (Benth.) Podlech & Iranshahr | - | KT031977 | - |
| *Nanorrhinum petranum* (Danin) Yousefi & Zarre | KT031909 | KT031968 | - |
| *Nanorrhinum ramosissimum* (Wall.) Betsche | KT031904 | KT031963 | - |
| *Nanorrhinum sagittatum* (Poir.) Yousefi & Zarre | KT031902 | KT031961 | - |
| *Nanorrhinum scariosepalum* (Täckh. & Boulos) Yousefi & Zarre | KT031911 | KT031970 | - |
| *Nanorrhinum scoparium* (Brouss. ex Spreng.) Yousefi & Zarre | KT031903 | KT031962 | - |
| *Nanorrhinum urbani* (Pit.) Yousefi & Zarre | KT031915 | KT031974 | - |
| *Nanorrhinum woodii* (D.A.Sutton) Ghebr. | MK095670 | MK095746 | - |
| *Neogaerrhinum filipes* (A.Gray) Rothm. | AF513896 | **forthcoming** | KT187845 |
| *Neogaerrhinum strictum* (Hook. & Arn.) Rothm. | AF513904 | **forthcoming** | JN848490 |
| *Pseudomisopates rivas-martinezii* (Sánchez Mata) Güemes | AY731262 | KM104750 | JN848492 |
| *Pseudorontium cyathiferum* (Benth.) Rothm. | AF513884 | - | JN848493 |
| *Rhodochiton atrosanguineum* (Zucc.) Rothm. | AF513876 | - | AJ250390 |
| *Sairocarpus breweri* (A.Gray) D.A.Sutton | KT187749 | - | KT187847 |
| *Sairocarpus cornutus* (Benth.) D.A.Sutton | AF513905 | **forthcoming** | KT187848 |
| *Sairocarpus cornutus* subsp. *leptaelus* (A.Gray) Barringer | **forthcoming** | **forthcoming** | - |
| *Sairocarpus costatus* (Wiggins) D.A.Sutton | AF513893 | - | KT187849 |
| *Sairocarpus coulterianus* (Benth. ex A.DC.) D.A.Sutton | KT187750 | **forthcoming** | KT187850 |
| *Sairocarpus elmeri* (Rothm.) D.A.Sutton | **forthcoming** | **forthcoming** | - |
| *Sairocarpus kingii* (S.Watson) D.A.Sutton | AF513903 | **forthcoming** | KT187851 |
| *Sairocarpus multiflorus* D.A.Sutton | AF513897 | **forthcoming** | KT187853 |
| *Sairocarpus nuttallianus* (Benth. ex A.DC.) D.A.Sutton | AF513895 | **forthcoming** | KT187852 |
| *Sairocarpus subcordatus* (A.Gray) D.A.Sutton | AF513902 | **forthcoming** | KT187854 |
| *Sairocarpus vexillocalyculatus* (Kellogg) D.A.Sutton | AF513900 | **forthcoming** | KT187855 |
| *Sairocarpus vexillocalyculatus* subsp. *intermedius* (D.M.Thomps.) Barringer | AF513907 | - | - |
| *Sairocarpus virga* (A.Gray) D.A.Sutton | AF513898 | **forthcoming** | KT187856 |
| *Sairocarpus watsonii* (Vasey & Rose) D.A.Sutton | AF513894 | - | - |
| *Schweinfurthia imbricata* A.G.Mill., M.Short & D.A.Sutton | AY731254 | **forthcoming** | KT187857 |
| *Schweinfurthia latifolia* Baker ex Oliver | AY731255 | **forthcoming** | KT187858 |
| *Schweinfurthia papilionacea* (L.) Boiss. | AY731253 | KP851131 | JN848495 |
| *Schweinfurthia pedicellata* Benth. & Hook.f. | AY731256 | **forthcoming** | KT187859 |
| *Schweinfurthia pterosperma* (A. Richard) A. Braun | AF513882 | - | - |
| *Schweinfurthia spinosa* A.G.Mill., M.Short & D.A.Sutton | AY731257 | **forthcoming** | KT187860 |

**Table S2.** Vouchers specimens for newly-sequenced taxa of Antirrhineae.

| Taxon | Voucher |
|---|---|
| **INGROUP (PLANTAGINACEAE, ANTIRRHINEAE)** | |
| *Anarrhinum corsicum* Jord. & Fourr | forthcoming |
| *Antirrhinum austral* Rothm. | 40IML11(2) |
| *Antirrhinum braun-blanquetii* Rothm. | 75PV07 |
| *Antirrhinum charidemi* Lange | 169PV05 |
| *Antirrhinum cirrhigerum* (Welw. ex Ficalho) Rothm. | 64PV10(1) |
| *Antirrhinum controversum* Pau | 1IML11(1) |
| *Antirrhinum grosii* Font Quer | 136PV10(1) |
| *Antirrhinum hispanicum* Chav. | 9IML13(2) |
| *Antirrhinum latifolium* Mill. | 25IML11(1) |
| *Antirrhinum linkianum* Boiss. & Reut. | forthcoming |
| *Antirrhinum litigiosum* Pau | 141PV08 |
| *Antirrhinum lopesianum* Rothm. | 10IML11(2) |
| *Antirrhinum meonanthum* Hoffmans. & Link | 33PV07 |
| *Antirrhinum microphyllum* Rothm. | 15PV08 |
| *Antirrhinum molle* L. | 83PV104 |
| *Antirrhinum mollissimum* (Pau) Rothm. | 51PV07 |
| *Antirrhinum onubense* (Fern.Casas) Fern.Casas | forthcoming |
| *Antirrhinum pertegasii* Pau ex Rothm. | J.Güemes |
| *Antirrhinum pulverulentum* Lázaro Ibiza | forthcoming |
| *Antirrhinum rothmaleri* (P.Silva) Amich, Bernardos & García-Barriuso | forthcoming |
| *Antirrhinum sempervirens* Lapeyr. | forthcoming |
| *Antirrhinum siculum* Mill. | Salvatore |
| *Antirrhinum subbaeticum* Güemes, Mateu & Sánchez-Gómez | MA |
| *Antirrhinum tortuosum* Bosc ex Vent. | VAL39871 |
| *Antirrhinum valentinum* Font Quer | VAL6, E.Carrio |
| *Galvezia ballii* Munz | 133PV07(13) |
| *Galvezia elisensii* M.O.Dillon & Quip. | forthcoming |
| *Galvezia lanceolata* Pennell | WS612 |
| *Galvezia leucantha* Wiggins | B.Sullivan 1-BS |
| *Howelliella ovata* (Eastw.) Rothm. | forthcoming |
| *Kickxia aegyptiaca* subsp. *fruticosa* (Desf.) Wickens | A00 & FJVS 137A00 |
| *Linaria brachyceras* (Bunge) Kuprian. | Aldasoro A23068 |
| *Linaria purpurea* (L.) Mill. | J. Ruiz s.n. |
| *Linaria sessilis* Kuprian. | Aldasoro A23028 |
| *Lophospermum breedlovei* Elisens | W.Elisens 713 |
| *Lophospermum erectum* (Hemsl.) Rothm. | GH2519 |
| *Lophospermum erubescens* D.Don | forthcoming |
| *Lophospermum purpusii* (Brandegee) Rothm. | forthcoming |
| *Lophospermum scandens* D.Don | D.Gordon 765 |
| *Lophospermum turneri* Elisens | W.Elisens 697 |
| *Mabrya acerifolia* (Pennell) Elisens | A.Forrest |
| *Mabrya coccinea* (I.M.Johnst.) Elisens | W.Elisens 562 |

| Taxon | Voucher |
|---|---|
| *Mabrya rosei* (Munz) Elisens | A.Forrest |
| *Maurandella antirrhiniflora* (Willd.) Rothm. | forthcoming |
| *Maurandya barclaiana* Lindl. | forthcoming |
| *Maurandya scandens* (Cav.) Pers. | forthcoming |
| *Misopates calycinum* (Vent.) Rothm. | 8MF08 bis |
| *Misopates microcarpum* (Pomel) D.A.Sutton | 13PV08 |
| *Neogaerrhinum filipes* (A.Gray) Rothm. | Thompson 254 |
| *Neogaerrhinum strictum* (Hook. & Arn.) Rothm. | A257572 |
| *Sairocarpus cornutus* (Benth.) D.A.Sutton | A257571 |
| *Sairocarpus cornutus* subsp. *leptaelus* (A.Gray) Barringer | forthcoming |
| *Sairocarpus coulterianus* (Benth. ex A.DC.) D.A.Sutton | 588893MA |
| *Sairocarpus elmeri* (Rothm.) D.A.Sutton | forthcoming |
| *Sairocarpus kingie* (S.Watson) D.A.Sutton | Morefield 3382 |
| *Sairocarpus multiflorus* D.A.Sutton | A269968 |
| *Sairocarpus nuttallianus* (Benth. ex A.DC.) D.A.Sutton | MA494669 |
| *Sairocarpus subcordatus* (A.Gray) D.A.Sutton | R.Oyama RK79 |
| *Sairocarpus vexillocalyculatus* (Kellogg) D.A.Sutton | R.Oyama #91 |
| *Sairocarpus virga* (A.Gray) D.A.Sutton | A269963 |
| *Schweinfurthia imbricata* A.G.Mill., M.Short & D.A.Sutton | E99215 |
| *Schweinfurthia latifolia* Baker ex Oliver | E99214 |
| *Schweinfurthia pedicellata* Benth. &Hook.f. | E99213 |
| *Schweinfurthia spinosa* A.G.Mill., M.Short & D.A.Sutton | E99203 |

**Table S3.** Distribution ranges of taxa used in biogeographic analyses. In the cases where subspecies are included, the range under the species name represents that of the type subspecies. Sources of information are provided. Areas: NE, Nearctic Region; WP, Western Palearctic Region; EP, Eastern Palearctic Region; MA, Madrean Region; ME, Mediterranean Region; IT, Irano-Turanian Region; NT, Neotropical Region; MC, Macaronesian Region; AF, African Region; IN, Indian-Indochinese Region.

| **Taxon** | **Range** | **Source** |
|---|---|---|
| *Lafuentea jeanpertiana* | ME | (Maire, 1921) |
| *Lafuentea rotundifolia* | ME | (Amich, 2009b) |
| *Anarrhinum bellidifolium* | WP - ME | (Amich, 2009a; Sutton, 1988) |
| *Anarrhinum duriminium* | WP – ME | (Amich, 2009a; Sutton, 1988) |
| *Anarrhinum longipedicellatum* | ME | (Amich, 2009a; Sutton, 1988) |
| *Anarrhinum laxiflorum* | ME | (Amich, 2009a; Sutton, 1988) |
| *Anarrhinum pedatum* | ME | (Sutton, 1988) |
| *Anarrhinum forskaohlii* | ME - AF - IT | (Sutton, 1988) |
| *Anarrhinum corsicum* | ME | (Sutton, 1988) |
| *Anarrhinum fruticosum* | ME | (Amich, 2009a; Sutton, 1988) |
| *Kickxia spuria* | WP | (Sutton, 1988) |
| *Kickxia spuria* subsp. *integrifolia* | WP - MC - ME - AF - IT | (Güemes, 2009d; Sutton, 1988) |
| *Kickxia lanigera* | MC - ME | (Sutton, 1988) |
| *Kickxia dentata* | ME | (Sutton, 1988; Yousefi, Zarre, & Heubl, 2016) |
| *Kickxia floribunda* | AF | (Sutton, 1988) |
| *Kickxia aegyptiaca* | ME - AF | (Sutton, 1988) |
| *Kickxia aegyptiaca* subsp. *fruticosa* | ME | (Sutton, 1988) |
| *Kickxia cirrhosa* | ME | (Güemes, 2009d; Sutton, 1988) |
| *Kickxia commutata* | WP - MC - ME | (Güemes, 2009d; Sutton, 1988) |
| *Kickxia elatine* | WP - MC - ME | (Güemes, 2009d; Sutton, 1988) |
| *Kickxia elatine* subsp. *crinita* | WP - ME - AF - IT | (Güemes, 2009d; Sutton, 1988) |
| *Kickxia caucasica* | WP | (Fernandes, 1972; Kupriyanova, 1997; Yousefi et al., 2016) |
| *Kickxia sieberi* | ME - AF | (Hayek & Markgraf, 1931; Yousefi et al., 2016) |
| *Nanorrhinum acerbianum* | AF | (Sutton, 1988) |
| *Nanorrhinum judaicum* | AF | (Sutton, 1988) |
| *Nanorrhinum macilentum* | AF | (Ghebrehiwet, 2000; Sutton, 1988) |
| *Nanorrhinum petranum* | AF | (Danin, 1991) |
| *Nanorrhinum cabulicum* | AF - IT | (Sutton, 1988) |
| *Nanorrhinum ramosissimum* | AF - IT - IN | (Ghebrehiwet, 2000; Sutton, 1988) |
| *Nanorrhinum ovatum* | AF - IT | (Ghebrehiwet, 2000; Sutton, 1988) |
| *Nanorrhinum scariosepalum* | AF | (Sutton, 1988) |
| *Nanorrhinum elegans* | MC | (Sutton, 1988) |
| *Nanorrhinum woodii* | AF | (Ghebrehiwet, 2000; Sutton, 1988) |
| *Nanorrhinum asparagoides* | AF | (Ghebrehiwet, 2000; Sutton, 1988) |

| Taxon | Range | Source |
|---|---|---|
| *Nanorrhinum heterophyllum* subsp. *canariensis* | MC | (Sutton, 1988) |
| *Nanorrhinum heterophyllum* | ME - AF | (Sutton, 1988) |
| *Nanorrhinum scoparium* | MC | (Sutton, 1988) |
| *Nanorrhinum urbani* | MC | (Sutton, 1988) |
| *Epixiphium wislizeni* | MA | (Sutton, 1988) |
| *Holmgrenanthe petrophila* | MA | (Elisens, 1985; Sutton, 1988) |
| *Maurandella antirrhiniflora* | NE - MA | (Elisens, 1985; Sutton, 1988) |
| *Maurandya barclaiana* | MA | (Elisens, 1985; Sutton, 1988) |
| *Lophospermum breedlovei* | NT | (Elisens, 1985; Sutton, 1988) |
| *Rhodochiton atrosanguineum* | MA | (Elisens, 1985; Sutton, 1988) |
| *Lophospermum erectum* | MA | (Sutton, 1988) |
| *Mabrya rosei* | MA | (Elisens, 1985; Sutton, 1988) |
| *Mabrya coccinea* | MA | (Elisens, 1985; Sutton, 1988) |
| *Mabrya acerifolia* | MA | (Elisens, 1985; Sutton, 1988) |
| *Mabrya geniculata* | MA | (Elisens, 1985; Sutton, 1988) |
| *Lophospermum turneri* | NT | (Elisens, 1985; Sutton, 1988) |
| *Lophospermum erubescens* | MA | (Elisens, 1985; Sutton, 1988) |
| *Maurandya scandens* | MA - NT | (Elisens, 1985; Sutton, 1988) |
| *Lophospermum purpusii* | MA | (Elisens, 1985; Sutton, 1988) |
| *Lophospermum scandens* | MA - NT | (Elisens, 1985; Sutton, 1988) |
| *Asarina procumbens* | WP | (Güemes, 2009b; Sutton, 1988) |
| *Gadoria falukei* | ME | (Güemes & Mota, 2017) |
| *Cymbalaria microcalyx* | ME | (Sutton, 1988) |
| *Cymbalaria aequitriloba* | ME | (Güemes, 2009c; Sutton, 1988) |
| *Cymbalaria hepaticifolia* | ME | (Sutton, 1988) |
| *Cymbalaria fragilis* | ME | (Güemes, 2009c; Sutton, 1988) |
| *Cymbalaria muelleri* | ME | (Sutton, 1988) |
| *Cymbalaria longipes* | ME | (Sutton, 1988) |
| *Cymbalaria microcalyx* subsp. *minor* | ME | (Sutton, 1988) |
| *Cymbalaria pubescens* | ME | (Sutton, 1988) |
| *Cymbalaria microcalyx* subsp. *acutiloba* | ME | (Sutton, 1988) |
| *Cymbalaria microcalyx* subsp. *dodekanesii* | ME | (Sutton, 1988) |
| *Cymbalaria muralis* | WP - ME | (Güemes, 2009c; Sutton, 1988) |
| *Cymbalaria glutinosa* | ME | (Bigazzi & Raffaelli, 2000; Sutton, 1988) |
| *Cymbalaria glutinosa* subsp. *brevicalcarata* | ME | (Bigazzi & Raffaelli, 2000; Sutton, 1988) |
| *Cymbalaria microcalyx* subsp. *ebelii* | ME | (Sutton, 1988) |
| *Cymbalaria muralis* subsp. *visianii* | ME | (Sutton, 1988) |
| *Cymbalaria pallida* | ME | (Sutton, 1988) |
| *Schweinfurthia pedicellata* | AF | (Sutton, 1988) |
| *Schweinfurthia pterosperma* | AF | (Sutton, 1988) |
| *Schweinfurthia imbricata* | AF | (Sutton, 1988) |
| *Schweinfurthia papilionacea* | AF - IT | (Sutton, 1988) |
| *Schweinfurthia latifolia* | AF | (Sutton, 1988) |
| *Schweinfurthia spinosa* | AF | (Sutton, 1988) |
| *Pseudorontium cyathiferum* | MA - NT | (Sutton, 1988) |

| Taxon | Range | Source |
|---|---|---|
| *Galvezia balli* | NT | (Sutton, 1988) |
| *Galvezia fruticosa* | NT | (Sutton, 1988) |
| *Galvezia elisensii* | NT | (Dillon & Quipuscoa Silvestre, 2014) |
| *Galvezia lanceolata* | NT | (Sutton, 1988) |
| *Galvezia leucantha* subsp. *pubescens* | NT | (Dillon & Quipuscoa Silvestre, 2014; Sutton, 1988) |
| *Galvezia leucantha* | NT | (Sutton, 1988) |
| *Galvezia leucantha* subsp. *porphyrantha* | NT | (Dillon & Quipuscoa Silvestre, 2014) |
| *Chaenorhinum cryptarum* | IT | (Sutton, 1988) |
| *Chaenorhinum pterosporum* | ME - IT | (Sutton, 1988) |
| *Chaenorhinum tenellum* | ME | (Benedí & Güemes, 2009; Sutton, 1988) |
| *Chaenorhinum litorale* | WP - ME | (Sutton, 1988) |
| *Chaenorhinum minus* | WP - ME | (Benedí & Güemes, 2009; Sutton, 1988) |
| *Chaenorhinum calycinum* | ME - AF - IT | (Sutton, 1988) |
| *Chaenorhinum huber-morathii* | IT | (Sutton, 1988) |
| *Albraunia foveopilosa* | IT | (Sutton, 1988) |
| *Chaenorhinum tuberculatum* | IT | (Sutton, 1988) |
| *Chaenorhinum johnstonii* | AF - IT | (Sutton, 1988) |
| *Chaenorrhinum spicatum* | IT | (Sutton, 1988) |
| *Chaenorhinum formenterae* | ME | (Benedí & Güemes, 2009; Sutton, 1988) |
| *Chaenorhinum crassifolium* | ME | (Benedí & Güemes, 2009; Sutton, 1988) |
| *Chaenorhinum rubrifolium* | ME | (Benedí & Güemes, 2009; Sutton, 1988) |
| *Chaenorhinum grandiflorum* | ME | (Benedí & Güemes, 2009; Sutton, 1988) |
| *Chaenorhinum glareosum* | ME | (Benedí & Güemes, 2009; Sutton, 1988) |
| *Chaenorhinum macropodum* | ME | (Benedí & Güemes, 2009; Sutton, 1988) |
| *Chaenorhinum raveyi* | ME | (Benedí & Güemes, 2009) |
| *Chaenorhinum reyesii* | ME | (Benedí & Güemes, 2009) |
| *Chaenorhinum robustum* | ME | (Benedí & Güemes, 2009; Sutton, 1988) |
| *Chaenorhinum villosum* | ME | (Benedí & Güemes, 2009; Sutton, 1988) |
| *Chaenorhinum flexuosum* | ME | (Sutton, 1988) |
| *Chaenorhinum exile* | ME | (Benedí & Güemes, 2009; Sutton, 1988) |
| *Chaenorhinum segoviense* | ME | (Benedí & Güemes, 2009; Sutton, 1988) |
| *Chaenorhinum origanifolium* | WP - ME | (Benedí & Güemes, 2009; Sutton, 1988) |
| *Chaenorhinum serpyllifolium* | ME | (Benedí & Güemes, 2009; Sutton, 1988) |
| *Acanthorrhinum ramosissimum* | ME - AF | (Sutton, 1988) |
| *Pseudomisopates rivas-martinezii* | ME | (Güemes,1997) |
| *Misopates microcarpum* | ME - AF | (Güemes, 2009e; Sutton, 1988) |
| *Misopates calycinum* | MC - ME | (Güemes, 2009e; Sutton, 1988) |
| *Misopates chrysothales* | ME | (Sutton, 1988) |
| *Misopates orontium* | WP - MC - ME - AF - IT - IN | (Güemes, 2009e; Sutton, 1988) |
| *Sairocarpus kingii* | MA | (Sutton, 1988) |
| *Mohavea breviflora* | MA | (Sutton, 1988) |
| *Mohavea confertiflora* | MA | (Sutton, 1988) |
| *Howelliella ovata* | MA | (Sutton, 1988) |
| *Sairocarpus breweri* | NE | (Barringer, 2013; Sutton, 1988) |
| *Sairocarpus elmeri* | MA | (Sutton, 1988) |

| **Taxon** | **Range** | **Source** |
|---|---|---|
| *Sairocarpus subcordatus* | NE | (Sutton, 1988) |
| *Sairocarpus vexillocalyculatus* | NE - MA | (Barringer, 2013; Sutton, 1988) |
| *Sairocarpus vexillocalyculatus* subsp. *intermedius* | NE | (Barringer, 2013) |
| *Sairocarpus cornutus* | NE-MA | (Barringer, 2013; Sutton, 1988) |
| *Sairocarpus cornutus* subsp. *leptaleus* | NE-MA | (Barringer, 2013) |
| *Gambelia juncea* | MA | (Sutton, 1988) |
| *Gambelia speciosa* | MA | (Sutton, 1988) |
| *Sairocarpus costatus* | MA | (Sutton, 1988) |
| *Neogaerrhinum strictum* | MA | (Sutton, 1988) |
| *Sairocarpus nuttallianus* | MA | (Sutton, 1988) |
| *Neogaerrhinum filipes* | MA | (Sutton, 1988) |
| *Sairocarpus coulterianus* | MA | (Sutton, 1988) |
| *Sairocarpus watsonii* | MA | (Sutton, 1988) |
| *Sairocarpus multiflorus* | MA | (Sutton, 1988) |
| *Sairocarpus virga* | NE | (Sutton, 1988) |
| *Antirrhinum controversum* | ME | (Güemes, 2009a; Sutton, 1988) |
| *Antirrhinum siculum* | ME | (Sutton, 1988) |
| *Antirrhinum valentinum* | ME | (Güemes, 2009a; Sutton, 1988) |
| *Antirrhinum australe* | ME | (Güemes, 2009a; Sutton, 1988) |
| *Antirrhinum subbaeticum* | ME | (Güemes, 2009a) |
| *Antirrhinum hispanicum* | ME | (Güemes, 2009a; Sutton, 1988) |
| *Antirrhinum charidemi* | ME | (Güemes, 2009a; Sutton, 1988) |
| *Antirrhinum mollissimum* | ME | (Güemes, 2009a; Sutton, 1988) |
| *Antirrhinum grosii* | ME | (Güemes, 2009a; Sutton, 1988) |
| *Antirrhinum meonanthum* | WP - ME | (Güemes, 2009a; Sutton, 1988) |
| *Antirrhinum onubense* | ME | (Güemes, 2009a) |
| *Antirrhinum microphyllum* | ME | (Güemes, 2009a; Sutton, 1988) |
| *Antirrhinum pulverulentum* | ME | (Güemes, 2009a; Sutton, 1988) |
| *Antirrhinum pertegasii* | ME | (Güemes, 2009a; Sutton, 1988) |
| *Antirrhinum linkianum* | WP - ME | (Güemes, 2009a; Sutton, 1988) |
| *Antirrhinum litigiosum* | WP - ME | (Güemes, 2009a; Sutton, 1988) |
| *Antirrhinum graniticum* | ME | (Güemes, 2009a; Sutton, 1988) |
| *Antirrhinum cirrhigerum* | ME | (Güemes, 2009a; Sutton, 1988) |
| *Antirrhinum tortuosum* | ME | (Güemes, 2009a; Sutton, 1988) |
| *Antirrhinum sempervirens* | WP | (Güemes, 2009a; Sutton, 1988) |
| *Antirrhinum rothmaleri* | ME | (García-Barriuso et al., 2011) |
| *Antirrhinum braun-blanquetii* | WP | (Güemes, 2009a; Sutton, 1988) |
| *Antirrhinum lopesianum* | ME | (Güemes, 2009a; Sutton, 1988) |
| *Antirrhinum latifolium* | WP - ME | (Güemes, 2009a; Sutton, 1988) |
| *Antirrhinum majus* | WP - ME | (Güemes, 2009a; Sutton, 1988) |
| *Antirrhinum molle* | WP | (Güemes, 2009a; Sutton, 1988) |
| *Linaria elegans* | WP - ME | (Sáez & Bernal, 2009; Sutton, 1988) |
| *Linaria nigricans* | ME | (Sáez & Bernal, 2009; Sutton, 1988) |
| *Linaria galioides* | ME | (Sutton, 1988) |
| *Linaria tingitana* | ME | (Sutton, 1988) |

| **Taxon** | **Range** | **Source** |
|---|---|---|
| *Linaria gattefossei* | ME | (Maire, 1938) |
| *Linaria bordiana* subsp. *kralikiana* | ME | (Sutton, 1988) |
| *Linaria imzica* | ME | (Gómiz, 2004) |
| *Linaria weilleri* | ME | (Sutton, 1988) |
| *Linaria viscosa* | ME | (Sáez & Bernal, 2009; Sutton, 1988) |
| *Linaria onubensis* | ME | (Vigalondo, Fernández-Mazuecos, Vargas, & Sáez, 2015) |
| *Linaria spartea* | WP - ME | (Sáez & Bernal, 2009; Sutton, 1988) |
| *Linaria algarviana* | ME | (Sáez & Bernal, 2009; Sutton, 1988) |
| *Linaria incarnata* | ME | (Sáez & Bernal, 2009; Sutton, 1988) |
| *Linaria becerrae* | ME | (Blanca, Cueto, & Fuentes, 2017) |
| *Linaria clementei* | ME | (Sáez & Bernal, 2009; Sutton, 1988) |
| *Linaria salzmannii* | ME | (Blanca et al., 2017) |
| *Linaria mamorensis* | ME | (Vigalondo et al., 2015) |
| *Linaria pedunculata* | ME | (Sáez & Bernal, 2009; Sutton, 1988) |
| *Linaria gharbensis* | ME | (Sáez & Bernal, 2009; Sutton, 1988) |
| *Linaria hellenica* | ME | (Sutton, 1988) |
| *Linaria bipartita* | ME | (Sutton, 1988) |
| *Linaria multicaulis* subsp. *heterophylla* | ME | (Sutton, 1988) |
| *Linaria maroccana* | ME | (Sutton, 1988) |
| *Linaria pinifolia* | ME | (Sutton, 1988) |
| *Linaria iranica* | IT | (Rahmani, Nejadsatari, Hamdi, Mehregan, & Assadi, 2014) |
| *Linaria tenuis* | ME - AF | (Sutton, 1988) |
| *Linaria bordiana* | ME | (Sutton, 1988) |
| *Linaria pseudoviscosa* | ME | (Sutton, 1988) |
| *Linaria multicaulis* | ME | (Sutton, 1988) |
| *Linaria multicaulis* subsp. *aurasiaca* | ME | (Sutton, 1988) |
| *Linaria pelisseriana* | WP - ME | (Sáez & Bernal, 2009; Sutton, 1988) |
| *Linaria triornithophora* | WP - ME | (Sáez & Bernal, 2009; Sutton, 1988) |
| *Linaria armeniaca* | WP - IT | (Rahmani et al., 2014; Sutton, 1988) |
| *Linaria chalepensis* | ME - AF - IT | (Rahmani et al., 2014; Sáez & Bernal, 2009; Sutton, 1988) |
| *Linaria texana* | NE - MA | (Crawford & Elisens, 2006; Sutton, 1988) |
| *Linaria canadense* | NE | (Crawford & Elisens, 2006; Sutton, 1988) |
| *Linaria subandina* | NT | (Sutton, 1988) |
| *Linaria hirta* | ME | (Sáez & Bernal, 2009; Sutton, 1988) |
| *Linaria orbensis* | ME | (Sáez & Bernal, 2009) |
| *Linaria faucicola* | WP | (Sáez & Bernal, 2009; Sutton, 1988) |
| *Linaria filicaulis* | WP | (Sáez & Bernal, 2009; Sutton, 1988) |
| *Linaria alpina* | WP - ME | (Sáez & Bernal, 2009; Sutton, 1988) |
| *Linaria arenaria* | WP | (Sutton, 1988) |
| *Linaria oligantha* | ME | (Sáez & Bernal, 2009; Sutton, 1988) |
| *Linaria ricardoi* | ME | (Sáez & Bernal, 2009; Sutton, 1988) |
| *Linaria bubanii* | WP | (Sáez & Bernal, 2009; Sutton, 1988) |
| *Linaria amethystea* subsp. *broussonetii* | ME | (Sutton, 1988) |
| *Linaria munbyana* | ME | (Sáez & Bernal, 2009; Sutton, 1988) |
| *Linaria propinqua* | WP - ME | (Sáez & Bernal, 2009; Sutton, 1988) |

| Taxon | Range | Source |
|---|---|---|
| *Linaria badalii* | WP - ME | (Sáez & Bernal, 2009; Sutton, 1988) |
| *Linaria glauca* | ME | (Sáez & Bernal, 2009; Sutton, 1988) |
| *Linaria glauca* subsp. *olcadium* | ME | (Sáez & Bernal, 2009; Sutton, 1988) |
| *Linaria huteri* | ME | (Sáez & Bernal, 2009; Sutton, 1988) |
| *Linaria intricata* | ME | (Sáez & Bernal, 2009; Sutton, 1988) |
| *Linaria bipunctata* | ME | (Sáez & Bernal, 2009; Sutton, 1988) |
| *Linaria ficalhoana* | ME | (Sáez & Bernal, 2009; Sutton, 1988) |
| *Linaria diffusa* | ME | (Sáez & Bernal, 2009; Sutton, 1988) |
| *Linaria amethystea* | WP - ME | (Sáez & Bernal, 2009; Sutton, 1988) |
| *Linaria saxatilis* | WP - ME | (Sáez & Bernal, 2009; Sutton, 1988) |
| *Linaria benitoi* | ME | (Sáez & Bernal, 2009) |
| *Linaria atlantica* | ME | (Sutton, 1988) |
| *Linaria tursica* | ME | (Sáez & Bernal, 2009; Sutton, 1988) |
| *Linaria arvensis* | WP - MC - ME - IT | (Rahmani et al., 2014; Sutton, 1988) |
| *Linaria micrantha* | MC - ME - AF - IT | (Reyes-Betancort, León, & Wildpret, 1999; Sutton, 1988) |
| *Linaria kavirensis* | IT | (Rahmani et al., 2014) |
| *Linaria simplex* | WP - MC - ME - AF - IT | (Rahmani et al., 2014; Sáez & Bernal, 2009; Sutton, 1988) |
| *Linaria nevadensis* | ME | (Sáez & Bernal, 2009; Sutton, 1988) |
| *Linaria tristis* subsp. *lurida* | ME | (Sutton, 1988) |
| *Linaria depauperata* | ME | (Sáez & Bernal, 2009) |
| *Linaria depauperata* subsp. *heglmaieri* | ME | (Sáez & Bernal, 2009) |
| *Linaria polygalifolia* subsp. *lamarckii* | ME | (Sáez & Bernal, 2009; Sutton, 1988) |
| *Linaria polygalifolia* | WP - ME | (Sáez & Bernal, 2009; Sutton, 1988) |
| *Linaria polygalifolia* subsp. *aguillonensis* | WP | (Sáez & Bernal, 2009) |
| *Linaria depauperata* subsp. *ilergabona* | ME | (Sáez & Bernal, 2009) |
| *Linaria atrofusca* | ME | (Sáez & Bernal, 2009) |
| *Linaria supina* | WP - ME | (Sáez & Bernal, 2009; Sutton, 1988) |
| *Linaria supina* subsp. *maritima* | WP | (Sáez & Bernal, 2009) |
| *Linaria aeruginea* | ME | (Sáez & Bernal, 2009; Sutton, 1988) |
| *Linaria tristis* subsp. *mesatlantica* | ME | (Sutton, 1988) |
| *Linaria tristis* subsp. *pectinata* | ME | (Sutton, 1988) |
| *Linaria lilacina* | ME | (Sutton, 1988) |
| *Linaria caesia* | ME | (Sáez & Bernal, 2009; Sutton, 1988) |
| *Linaria thymifolia* | WP | (Sutton, 1988) |
| *Linaria almijarensis* | ME | (Sáez & Bernal, 2009) |
| *Linaria anticaria* | ME | (Sutton, 1988) |
| *Linaria tristis* | ME | (Sáez & Bernal, 2009; Sutton, 1988) |
| *Linaria verticiliata* subsp. *cuartanensis* | ME | (Sáez & Bernal, 2009) |
| *Linaria saturejoides* | ME | (Sáez & Bernal, 2009; Sutton, 1988) |
| *Linaria oblongifolia* | ME | (Sáez & Bernal, 2009; Sutton, 1988) |
| *Linaria accitensis* | ME | (Sáez & Bernal, 2009) |
| *Linaria oblongifolia* subsp. *haenseleri* | ME | (Sáez & Bernal, 2009; Sutton, 1988) |
| *Linaria platycalyx* | ME | (Sáez & Bernal, 2009; Sutton, 1988) |
| *Linaria verticillata* | ME | (Sáez & Bernal, 2009; Sutton, 1988) |
| *Linaria amoi* | ME | (Sáez & Bernal, 2009; Sutton, 1988) |

| Taxon | Range | Source |
|---|---|---|
| *Linaria glacialis* | ME | (Sáez & Bernal, 2009; Sutton, 1988) |
| *Linaria latifolia* | ME | (Sáez & Bernal, 2009; Sutton, 1988) |
| *Linaria haelava* | ME - AF | (Sutton, 1988) |
| *Linaria joppensis* | ME | (Sutton, 1988) |
| *Linaria laxiflora* | ME | (Sutton, 1988) |
| *Linaria peltieri* | ME | (Sutton, 1988) |
| *Linaria warionis* | ME | (Sutton, 1988) |
| *Linaria decipiens* | ME | (Sutton, 1988) |
| *Linaria reflexa* | ME | (Sutton, 1988) |
| *Linaria cavanillesii* | ME | (Sáez & Bernal, 2009; Sutton, 1988) |
| *Linaria nivea* | ME | (Sáez & Bernal, 2009; Sutton, 1988) |
| *Linaria purpurea* | ME | (Sutton, 1988) |
| *Linaria capraria* | ME | (Sutton, 1988) |
| *Linaria repens* | WP - ME | (Sáez & Bernal, 2009; Sutton, 1988) |
| *Linaria ventricosa* | ME | (Sutton, 1988) |
| *Linaria virgata* | ME | (Sutton, 1988) |
| *Linaria parviracemosa* | ME | (Sutton, 1988) |
| *Linaria triphylla* | ME | (Sáez & Bernal, 2009; Sutton, 1988) |
| *Linaria flava* | ME | (Sutton, 1988) |
| *Linaria albifrons* | WP - ME - AF - IT | (Rahmani et al., 2014; Sutton, 1988) |
| *Linaria boushehrensis* | AF | (Rahmani et al., 2014) |
| *Linaria microsepala* | ME | (Sutton, 1988) |
| *Linaria peloponnesiaca* | ME | (Sutton, 1988) |
| *Linaria corifolia* | ME - IT | (Sutton, 1988) |
| *Linaria antilibanotica* | ME - IT | (Sutton, 1988) |
| *Linaria iconia* | ME - IT | (Sutton, 1988) |
| *Linaria orientalis* | IT | (Rahmani et al., 2014) |
| *Linaria golestanensis* | IT | (Rahmani et al., 2014) |
| *Linaria mazandaranensis* | IT | (Rahmani et al., 2014) |
| *Linaria unaiensis* | IT | (Sutton, 1988) |
| *Linaria bamianica* | IT | (Sutton, 1988) |
| *Linaria khorasanensis* | IT | (Rahmani et al., 2014) |
| *Linaria griffithsii* | IT | (Sutton, 1988) |
| *Linaria grandiflora* | WP - ME - IT | (Rahmani et al., 2014; Sutton, 1988) |
| *Linaria dalmatica* | WP - ME - IT | (Rahmani et al., 2014; Sutton, 1988) |
| *Linaria rubioides* | WP | (Sutton, 1988) |
| *Linaria shahroudensis* | IT | (Rahmani et al., 2014) |
| *Linaria transiliensis* | IT | (Sutton, 1988) |
| *Linaria incompleta* | WP - IT | (Sutton, 1988) |
| *Linaria cretacea* | WP | (Sutton, 1988) |
| *Linaria creticola* | WP | (Sutton, 1988) |
| *Linaria meyeri* | WP | (Sutton, 1988) |
| *Linaria pyramidalis* | IT | (Sutton, 1988) |
| *Linaria schelkownikowii* | WP - IT | (Sutton, 1988) |
| *Linaria leptoceras* | IT | (Sutton, 1988) |

| **Taxon** | **Range** | **Source** |
|---|---|---|
| *Linaria schirvanica* | IT | (Sutton, 1988) |
| *Linaria loeselii* | WP | (Sutton, 1988) |
| *Linaria odora* | WP - IT | (Rahmani et al., 2014; Sutton, 1988) |
| *Linaria brachyceras* | EP - IT | (Sutton, 1988) |
| *Linaria baligaliensis* | IT | (Sutton, 1988) |
| *Linaria sessilis* | IT | (Sutton, 1988) |
| *Linaria venosa* | IT | (Sutton, 1988) |
| *Linaria macroura* | WP | (Sutton, 1988) |
| *Linaria popovii* | IT | (Sutton, 1988) |
| *Linaria elymaitica* | IT | (Rahmani et al., 2014; Sutton, 1988) |
| *Linaria thibetica* | IT | (Sutton, 1988) |
| *Linaria japonica* | EP | (Sutton, 1988) |
| *Linaria altaica* | IT | (Sutton, 1988) |
| *Linaria vulgaris* | WP - EP - ME - IT | (Sáez & Bernal, 2009; Sutton, 1988) |
| *Linaria damascena* | ME | (Sutton, 1988) |
| *Linaria remotiflora* | IT | (Rahmani et al., 2014; Sutton, 1988) |
| *Linaria farsensis* | IT | (Rahmani et al., 2014) |
| *Linaria michauxii* | IT | (Rahmani et al., 2014; Sutton, 1988) |
| *Linaria kurdica* subsp. *pycnophylla* | IT | (Sutton, 1988) |
| *Linaria guilanensis* | IT | (Rahmani et al., 2014) |
| *Linaria kurdica* | WP - IT | (Podlech & Iranshahr, 2015; Sutton, 1988) |
| *Linaria angustissima* | WP - EP - ME | (Sutton, 1988) |
| *Linaria azerbaijanensis* | IT | (Rahmani et al., 2014) |
| *Linaria fastigiata* | IT | (Podlech & Iranshahr, 2015; Sutton, 1988) |
| *Linaria khalkhalensis* | IT | (Rahmani et al., 2014) |
| *Linaria confertiflora* | IT | (Podlech & Iranshahr, 2015; Sutton, 1988) |
| *Linaria karajensis* | IT | (Rahmani et al., 2014) |
| *Linaria genistifolia* | WP - ME - IT | (Sutton, 1988) |
| *Linaria pyramidalis* subsp. *kopetdaghensis* | IT | (Sutton, 1988) |
| *Linaria nurensis* | IT | (Rahmani et al., 2014; Sutton, 1988) |
| *Linaria lineolata* | IT | (Rahmani et al., 2014; Sutton, 1988) |
| *Linaria striatella* | IT | (Rahmani et al., 2014; Sutton, 1988) |

**Table S4.** Comparison of biogeographic models in BioGeoBEARS. Log-likelihood (LnL) and Akaike information criterion (AIC) are shown for each model. The best model is shown in bold.

| **Model** | **LnL** | **AIC** | **ΔAIC** |
|---|---|---|---|
| DEC | **-560.4** | **1124.8** | **0.0** |
| DIVALIKE | -597.0 | 1198.0 | 73.3 |
| BAYAREALIKE | -618.3 | 1240.7 | 115.9 |

**Fig. S1.** Dispersal probability matrices for each time slice (TS), and maps representing a schematic configuration of landmasses for each TS (based on Meseguer et al., 2015; see Appendix S1 for further information about dispersal probability values and time slices). Areas: NE, Nearctic Region; WP, Western Palearctic Region; EP, Eastern Palearctic Region; MA, Madrean Region; ME, Mediterranean Region; IT, Irano-Turanian Region; NT, Neotropical Region; MC, Macaronesian Region; AF, African Region; IN, Indian-Indochinese Region. Time Slices: TSI, 60-35 Ma; TSII, 35-10 Ma; TSIII, 10-3.5 Ma; TSIV, 3.5-0 Ma.

### Time Slice I: 60 - 35 Ma

| | WP | EP | NE | MC | ME | AF | IT | MA | IN | NT |
|---|---|---|---|---|---|---|---|---|---|---|
| WP | 1 | 0.5 | 1 | 0.5 | 1 | 0.1 | 0 | 0.1 | 0.1 | 0.1 |
| EP | 0.5 | 1 | 1 | 0.1 | 0.1 | 0.1 | 0 | 0.1 | 0.1 | 0.1 |
| NE | 1 | 1 | 1 | 0.1 | 0.5 | 0.1 | 0 | 1 | 0.1 | 0.1 |
| MC | 0.5 | 0.1 | 0.1 | 1 | 0.5 | 0.5 | 0 | 0.1 | 0.1 | 0.1 |
| ME | 1 | 0.1 | 0.5 | 0.5 | 1 | 1 | 0 | 0.1 | 0.1 | 0.1 |
| AF | 0.1 | 0.1 | 0.1 | 0.5 | 1 | 1 | 0 | 0.1 | 0.1 | 0.1 |
| IT | 0 | 0 | 0 | 0 | 0 | 0 | 1 | 0 | 0 | 0 |
| MA | 0.1 | 0.1 | 1 | 0.1 | 0.1 | 0.1 | 0 | 1 | 0.1 | 0.5 |
| IN | 0.1 | 0.1 | 0.1 | 0.1 | 0.1 | 0.1 | 0 | 0.1 | 1 | 0.1 |
| NT | 0.1 | 0.1 | 0.1 | 0.1 | 0.1 | 0.1 | 0 | 0.5 | 0.1 | 1 |

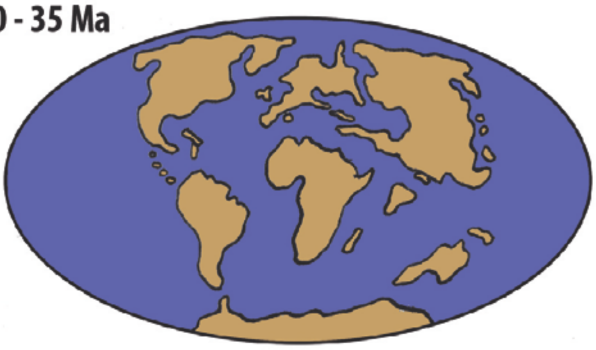

### Time Slice II: 35 - 10 Ma

| | WP | EP | NE | MC | ME | AF | IT | MA | IN | NT |
|---|---|---|---|---|---|---|---|---|---|---|
| WP | 1 | 1 | 0.5 | 0.5 | 1 | 0.1 | 0.5 | 0.1 | 0.1 | 0.1 |
| EP | 1 | 1 | 1 | 0.1 | 0.1 | 0.1 | 0.5 | 0.1 | 0.5 | 0.1 |
| NE | 0.5 | 1 | 1 | 0.1 | 0.1 | 0.1 | 0.1 | 1 | 0.1 | 0.1 |
| MC | 0.5 | 0.1 | 0.1 | 1 | 0.5 | 0.5 | 0.1 | 0.1 | 0.1 | 0.1 |
| ME | 1 | 0.1 | 0.1 | 0.5 | 1 | 1 | 0.5 | 0.1 | 0.1 | 0.1 |
| AF | 0.1 | 0.1 | 0.1 | 0.5 | 1 | 1 | 0.5 | 0.1 | 0.1 | 0.1 |
| IT | 0.5 | 0.5 | 0.1 | 0.1 | 0.5 | 0.5 | 1 | 0.1 | 0.5 | 0.1 |
| MA | 0.1 | 0.1 | 1 | 0.1 | 0.1 | 0.1 | 0.1 | 1 | 0.1 | 0.5 |
| IN | 0.1 | 0.5 | 0.1 | 0.1 | 0.1 | 0.1 | 0.5 | 0.1 | 1 | 0.1 |
| NT | 0.1 | 0.1 | 0.1 | 0.1 | 0.1 | 0.1 | 0.1 | 0.5 | 0.1 | 1 |

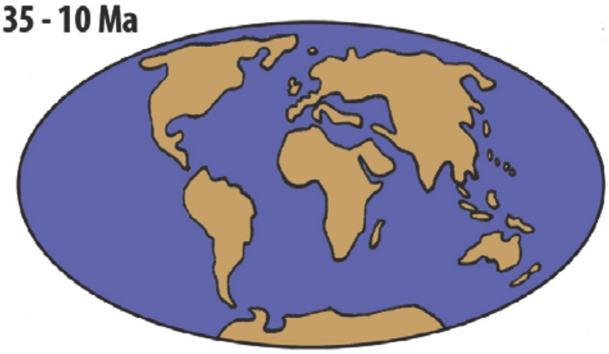

### Time Slice III: 10 - 3.5 Ma

| | WP | EP | NE | MC | ME | AF | IT | MA | IN | NT |
|---|---|---|---|---|---|---|---|---|---|---|
| WP | 1 | 1 | 0.1 | 0.5 | 1 | 0.1 | 1 | 0.1 | 0.1 | 0.1 |
| EP | 1 | 1 | 0.5 | 0.1 | 0.1 | 0.1 | 1 | 0.1 | 1 | 0.1 |
| NE | 0.1 | 0.5 | 1 | 0.1 | 0.1 | 0.1 | 0.1 | 1 | 0.1 | 0.5 |
| MC | 0.5 | 0.1 | 0.1 | 1 | 0.5 | 0.5 | 0.1 | 0.1 | 0.1 | 0.1 |
| ME | 1 | 0.1 | 0.1 | 0.5 | 1 | 1 | 1 | 0.1 | 0.1 | 0.1 |
| AF | 0.1 | 0.1 | 0.1 | 0.5 | 1 | 1 | 1 | 0.1 | 0.5 | 0.1 |
| IT | 1 | 1 | 0.1 | 0.1 | 1 | 1 | 1 | 0.1 | 1 | 0.1 |
| MA | 0.1 | 0.1 | 1 | 0.1 | 0.1 | 0.1 | 0.1 | 1 | 0.1 | 1 |
| IN | 0.1 | 1 | 0.1 | 0.1 | 0.1 | 0.5 | 1 | 0.1 | 1 | 0.1 |
| NT | 0.1 | 0.1 | 0.5 | 0.1 | 0.1 | 0.1 | 0.1 | 1 | 0.1 | 1 |

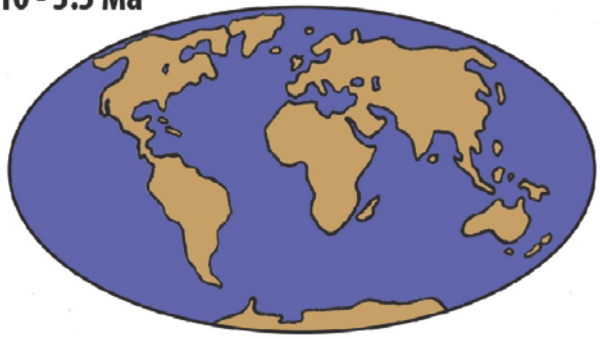

### Time Slice IV: 3.5 - 0 Ma

| | WP | EP | NE | MC | ME | AF | IT | MA | IN | NT |
|---|---|---|---|---|---|---|---|---|---|---|
| WP | 1 | 1 | 0.1 | 0.5 | 1 | 0.1 | 1 | 0.1 | 0.1 | 0.1 |
| EP | 1 | 1 | 0.1 | 0.1 | 0.1 | 0.1 | 1 | 0.1 | 1 | 0.1 |
| NE | 0.1 | 0.1 | 1 | 0.1 | 0.1 | 0.1 | 0.1 | 1 | 0.1 | 0.5 |
| MC | 0.5 | 0.1 | 0.1 | 1 | 0.5 | 0.5 | 0.1 | 0.1 | 0.1 | 0.1 |
| ME | 1 | 0.1 | 0.1 | 0.5 | 1 | 1 | 1 | 0.1 | 0.1 | 0.1 |
| AF | 0.1 | 0.1 | 0.1 | 0.5 | 1 | 1 | 1 | 0.1 | 0.5 | 0.1 |
| IT | 1 | 1 | 0.1 | 0.1 | 1 | 1 | 1 | 0.1 | 1 | 0.1 |
| MA | 0.1 | 0.1 | 1 | 0.1 | 0.1 | 0.1 | 0.1 | 1 | 0.1 | 1 |
| IN | 0.1 | 1 | 0.1 | 0.1 | 0.1 | 0.5 | 1 | 0.1 | 1 | 0.1 |
| NT | 0.1 | 0.1 | 0.5 | 0.1 | 0.1 | 0.1 | 0.1 | 1 | 0.1 | 1 |

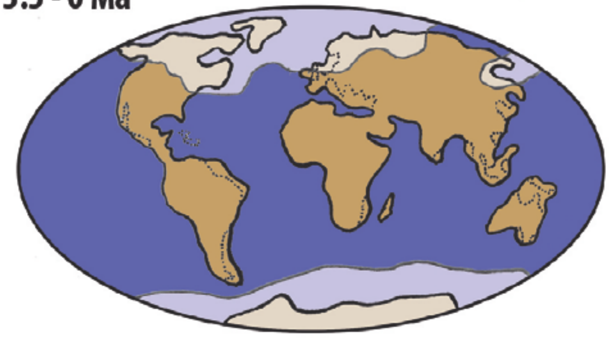

**Fig. S2.** Bayesian phylogenetic tree of Antirrhineae based on analysis of ITS, *ndh*F and *rpl*32-*trn*L sequences in MrBayes. Numbers above branches are Bayesian posterior probabilities. Node letters (A to M) indicate major clades within the Antirrhineae and the sister genus *Lafuentea* described in Appendix S1. The most recent common ancestors for the Plantaginaceae and the Antirrhineae are indicated and family names for outgroup taxa used for calibration are included.

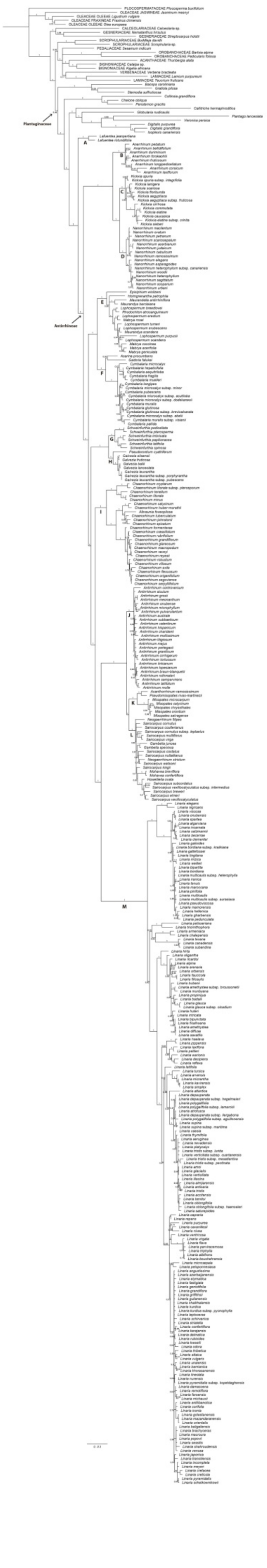
Plantaginaceae
Antirrhineae
A
B
C
D
E
F
G
H
I
J
K
L
M

**Fig. S3.** Maximum clade credibility tree from phylogenetic analysis in BEAST showing the Posterior Probability value for node support, blue bars indicate divergence time estimates at a 95% High Posterior Density. All other conventions as in Fig. S2.

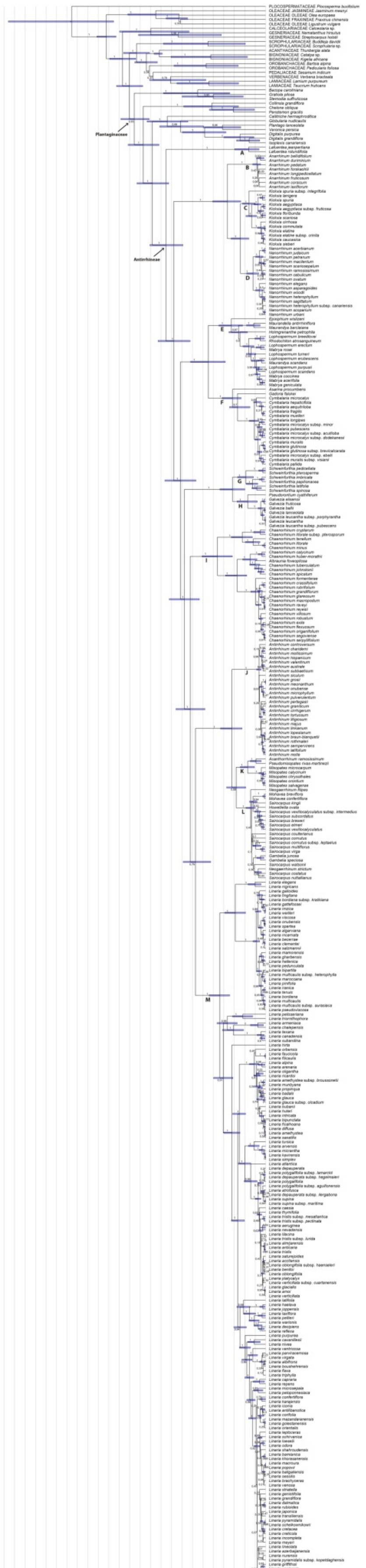

Plantaginaceae
Antirrhineae
A
B
C
D
E
F
G
H
I
J
K
L
M
125
100
75
50
25
0
(Ma)

**Fig. S4.** S-DIVA ancestral range estimation in RASP. Node letters (A to M) correspond to the major clades within Anitrrhineae and the sister genus *Lafuentea* described in Appendix S1. The most recent common ancestor for the Antirrhineae is indicated. The coloured range in the map represents the approximate distribution of the Antirrhineae. Areas: NE, Nearctic Region; WP, Western Palearctic Region; EP, Eastern Palearctic Region; MA, Madrean Region; ME, Mediterranean Region; IT, Irano-Turanian Region; NT, Neotropical Region; MC, Macaronesian Region; AF, African Region; IN, Indian-Indochinese Region.

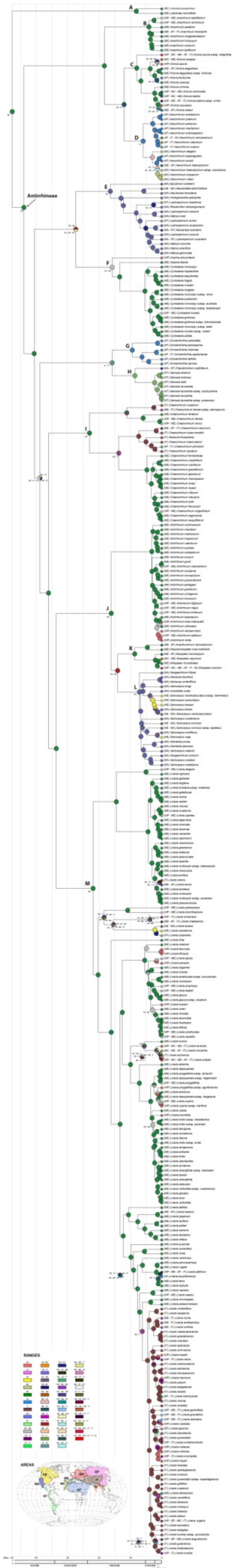

Antirrhineae
A
B
C
D
E
F
G
H
I
J
K
L
M
RANGES
AREAS

**Fig. S5.** Ancestral range estimation under the DEC model in BioGeoBEARS (with dispersal scalars). Letters at nodes and branches correspond to the range with the highest probability. Areas: A, Western Palearctic Region; B, Eastern Palearctic Region; C, Nearctic Region; D, Madrean Region; E, Mediterranean Region; F, African Region; G, Irano-Turanian Region; H, Macaronesian Region; I, Indian-Indochinese Region; J, Neotropical Region.

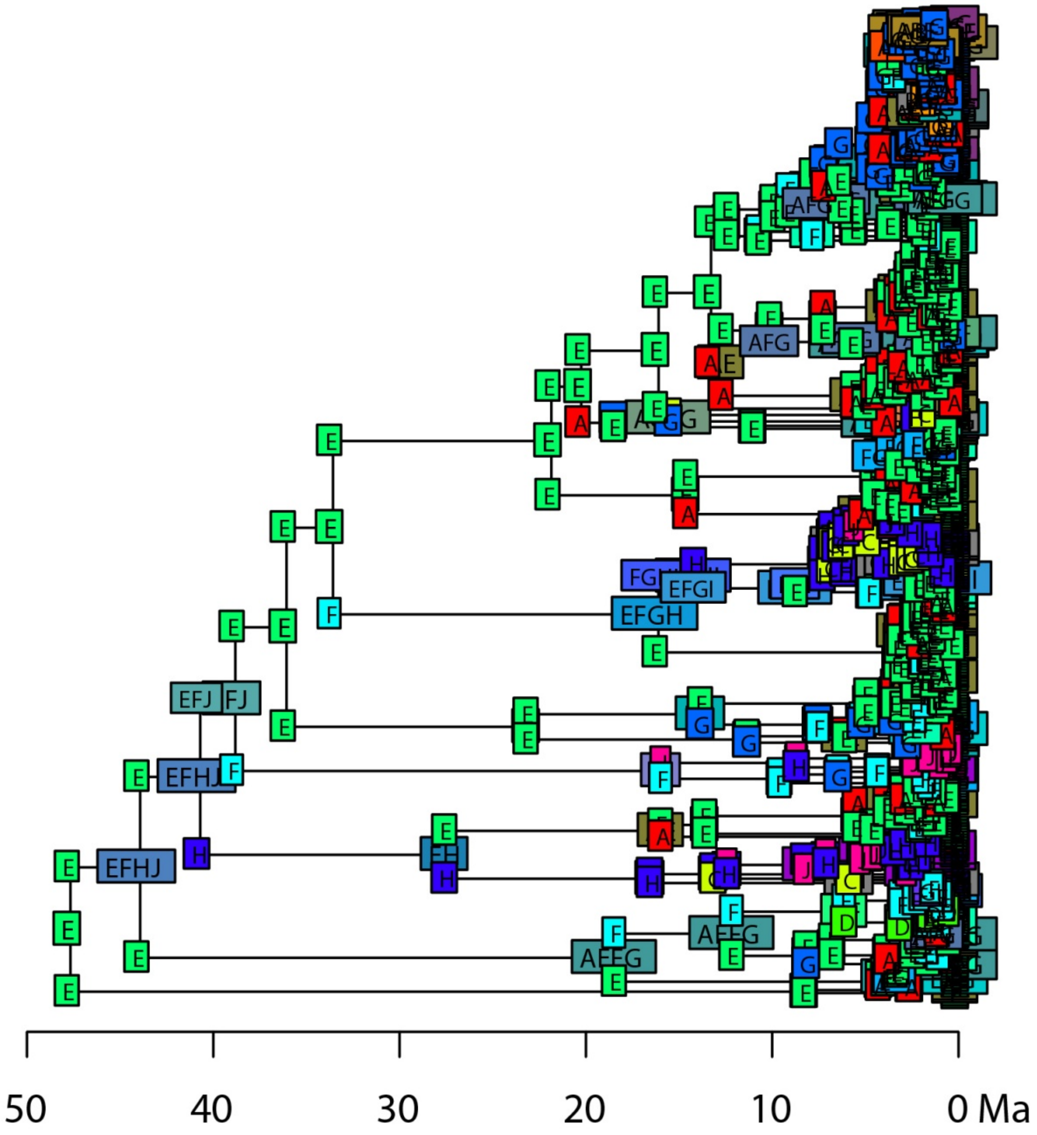

50
40
30
20
10
0 Ma
